# Rapid breath analysis for acute respiratory distress syndrome diagnostics using a portable 2-dimensional gas chromatography device

**DOI:** 10.1101/560888

**Authors:** Menglian Zhou, Ruchi Sharma, Hongbo Zhu, Jiliang Li, Shiyu Wang, Erin Bisco, Justin Massey, Amanda Pennington, Michael Sjoding, Robert P. Dickson, Pauline Park, Robert Hyzy, Lena Napolitano, Kevin R. Ward, Xudong Fan

## Abstract

Acute respiratory distress syndrome (ARDS) is the most severe form of acute lung injury, responsible for high mortality and long-term morbidity. As a dynamic syndrome with multiple etiologies its timely diagnosis is difficult as is tracking the course of the syndrome. Therefore, there is a significant need for early, rapid detection and diagnosis as well as clinical trajectory monitoring of ARDS. Here we report our work on using human breath to differentiate ARDS and non-ARDS causes of respiratory failure. A fully automated portable 2-dimensional gas chromatography device with high peak capacity, high sensitivity, and rapid analysis capability was designed and made in-house for on-site analysis of patients’ breath. A total of 85 breath samples from 48 ARDS patients and controls were collected. Ninety-seven elution peaks were separated and detected in 13 minutes. An algorithm based on machine learning, principal component analysis (PCA), and linear discriminant analysis (LDA) was developed. As compared to the adjudications done by physicians based on the Berlin criteria, our device and algorithm achieved an overall accuracy of 87.1% with 94.1% positive predictive value and 82.4% negative predictive value. The high overall accuracy and high positive predicative value suggest that the breath analysis method can accurately diagnose ARDS. The ability to continuously and non-invasively monitor exhaled breath for early diagnosis, disease trajectory tracking, and outcome prediction monitoring of ARDS may have a significant impact on changing practice and improving patient outcomes.

## 1. Introduction

Acute respiratory distress syndrome (ARDS) is an inflammatory condition of the lung producing severe lung damage. It is one of the most severe forms of acute lung injury and responsible for high mortality (40%) and long-term morbidity^1–3^. An estimated 200,000 Americans develop ARDS each year, of which more than 74,000 die from the disease^1^. Patients who survive ARDS experience long-term deficits in physical and neurocognitive function^4, 5^. Both primary hospitalizations and increased health service utilization among survivors are associated with high healthcare costs^1, 4^. For example, the average cost of an ICU (Intensive Care Unit) patient requiring mechanical ventilation ranges between $7,000 and $11,000 per day with an incremental cost of $1,000-1,500 per day for mechanical ventilation^6^.

Numerous predisposing factors for ARDS have previously been identified (*e.g*., sepsis, aspiration, and trauma)^7^. However, our understanding of patient susceptibility to ARDS is incomplete and the disease onset is poorly predicted by current risk models. Among patients with multiple established risk factors, the majority do not develop ARDS, while a minority develop severe, life-threatening disease^8, 9^. The most commonly used ARDS risk model (Lung Injury Prediction Score, LIPS) has strong negative predictive value (97%), effectively identifying patients at low risk for ARDS, but weak positive predictive value (18%),^1, 8^ implying poor ability to predict disease onset. The clinical diagnosis of ARDS is established based on the radiological, physiological, and clinical criteria summarized in the Berlin definition (Table 1)^9^. However, those criteria only show a moderate correlation with real time and post-mortem tissue pathological findings^10, 11^ and temporally lag the acute, dynamic inflammatory processes responsible for ARDS, and thus cannot be used for early diagnosis and trajectory monitoring of ARDS. Therefore, there is a significant unmet clinical need for early, rapid detection and diagnosis as well as clinical trajectory monitoring of ARDS.

**Table 1.** The Berlin Definition of Acute Respiratory Distress Syndrome (ARDS).

| **Acute Respiratory Distress Syndrome** |  |
| --- | --- |
| Timing | Within 1 week of a known clinical insult or new worsening respiratory symptoms |
| Chest imaging (Chest radiograph) | Bilateral opacities – not fully explained by effusion, lobar/lung collapse, or nodules |
| Origin of edema | Respiratory failure not fully explained by cardiac failure or fluid overload. Need objective assessment (e.g., Echocardiography) to exclude hydrostatic edema if no risk factor is present. |
| Oxygenation |  |
| Mild | $200 \text{ mmHg} < \text{PaO}_2 / \text{FiO}_2 \leq 300 \text{ mm Hg}$ with PEEP or CPAP $\geq 5 \text{ cm H}_2\text{O}$ |
| Moderate | $100 \text{ mmHg} < \text{PaO}_2 / \text{FiO}_2 \leq 200 \text{ mm Hg}$ with PEEP $\geq 5 \text{ cm H}_2\text{O}$ |
| Severe | $\text{PaO}_2 / \text{FiO}_2 \leq 100 \text{ mm Hg}$ with PEEP $\geq 5 \text{ cm H}_2\text{O}$ |
| Abbreviation: CPAP, continuous positive airway pressure; FiO<sub>2</sub>, fraction of inspired oxygen; PaO<sub>2</sub>, partial pressure of arterial oxygen; PEEP, positive end-expiratory pressure. |  |

Exhaled breath contains hundreds of volatile organic compounds (VOCs). Many VOCs (such as pentane, isoprene, and ethane) are related to inflammatory processes occurring in the lungs and systemically in blood from remote organ injury^12–15^. Those and other VOCs could potentially be used as biomarkers to predict the onset and severity of certain critical lung diseases such as ARDS as well as systemic inflammation such as sepsis. They also could help guide therapy if they could be measured simultaneously and precisely in real-time^16–19^. Unlike blood-based analysis, breath is unlimited in its sampling potential and can be noninvasively and continuously collected and analyzed. Technologies designed for the real-time analysis of VOCs in a point-of-care (POC) fashion could allow for the identification of breathomic signatures that enable the early diagnosis of ARDS, stratification, and trajectory monitoring, allowing for precision treatments.

Generally, there are a few technologies that can be used in breath analysis. GC-MS (gas chromatography in tandem with mass spectrometer) is the gold standard for the analysis of complex vapor mixtures such as breath samples. However, GC-MS is bulky, power intensive, and requires dedicated trained personnel. In practice, breath from a subject is collected in a sealed tube or bag, which is then sent to an analytical lab equipped with GC-MS for analysis. This method, of course, is not suitable for POC applications. In addition, due to long turn-around times, GC-MS cannot be used to continuously monitor the subject to detect dynamic changes. SIFT-MS (selected-ion flow-tube mass spectrometry) is another technology developed for breath analysis^20–22^. However, the bulky size, heavy weight (>200 kg), and high cost limit its wide acceptance. The third technology for breath analysis is based on ion mobility spectrometry (IMS)^23–25^. It can be operated under ambient pressure, thus avoiding the use of a cumbersome vacuum pump, which makes IMS portable. In addition, IMS based analysis is usually rapid. One assay typically takes just a few minutes. Recent exploratory tests using FAIMS (Field Asymmetric Ion Mobility Spectrometry) technology in diagnosis of lung cancer, asthma, and inflammatory bowel disease have been reported^24, 25^. Generally, when IMS is used in breath analysis, statistical and machine learning algorithm is employed to differentiate between target patients and healthy controls^24^. One of the major drawbacks of IMS, however, is its limited VOC separation capability, which may affect its diagnostic accuracy using breath as a target. Portable GC systems are the fourth technology for POC breath analysis. However, current commercial portable GC systems are 1-dimensional (1D) devices and have limited separation capability (or peak capacity), which, again, may affect the diagnostic accuracy. In addition, most of the 1D GC devices are not customized to operate in a fully automated mode and thus are not suitable for the use on a mechanical ventilator. Comprehensive 2D GC is a technique that is developed to improve the peak capacity over the traditional 1D GC.^26, 27^ In 2D GC, VOC analytes are subject to two independent separation processes, first by their vapor pressures in the 1^st^-dimensional column and then by their polarities in the 2^nd^-dimensional column. A 2D chromatogram can be constructed that consists of the 1^st^- and 2^nd^-dimensional retention times. Breath analysis with a commercial 2D GC instrument has been used for detection of diseases such as cancer, tuberculosis and human volatome^28–30^. Unfortunately, the existing commercial 2D GC systems are very bulky and expensive and require trained personnel. Therefore, they are not suitable for POC breath analysis and continuous monitoring. The fifth technology, e-Nose is developed as an indirect way for breath analysis, which relies on a vapor sensor array and pattern recognition^18, 31, 32^. For example, a colorimetric array device uses different dyes that respond collectively to distinguish various conditions from breath (through pattern recognition) among patients. While simple, e-Nose has poor chemical selectivity, device-to-device repeatability, and stability, as well as high susceptibility to background or interference VOCs^18, 19^. Very recently, a new technology was developed to collect and analyze the breath from mechanically ventilated patients using an in-line heat-moisture exchanger installed on a ventilator^33^. While potentially useful in ARDS diagnosis, this technology is focused on proteomic analysis of the breath condensates and requires long analysis time.

Recently we have developed a fully automated portable GC device that can be operated simultaneously as 1D GC and comprehensive 2D GC with a sub-ppb sensitivity^34^. With the help of the 2-dimensional separation, the co-eluted peaks that are not separated from the 1^st^-dimensional column can further be separated on the 2^nd^-dimensional column, thus increasing device’s separation capability. The aim of this study was to further adapt this portable GC for the use on a mechanical ventilator in ICUs and develop the related algorithms for rapid analysis of breath from patients undergoing mechanical ventilation, in order to understand the ability of our GC (and the algorithms) to detect the presence of ARDS compared to clinician adjudication.

Figure 1 shows the schematic of the GC device connected to a ventilator. In our work breath was collected and analyzed every 33 minutes via a small tube connected to the exhalation port of the ventilator. A total of 97 peaks were separated out from human breath. Through machine learning, principal component analysis (PCA), and linear discriminant analysis (LDA), 9 out of 97 peaks were selected as a VOC subset for the discrimination between ARDS and non-ARDS respiratory failure. Forty-eight (48) ARDS and non-ARDS patients with a total of 85 different breath chromatograms were evaluated. Among all 48 patients, we used 28 patients (43 sets of breath) as the training set and 20 patients (42 sets of breath) as the testing set. Using blinded physician adjudication of the patients’ records based on the Berlin criteria as the gold standard, our breath analysis achieved an overall accuracy of 87.1% with 94.1% positive predictive value and 82.4% negative predictive value.

**Figure 1.**
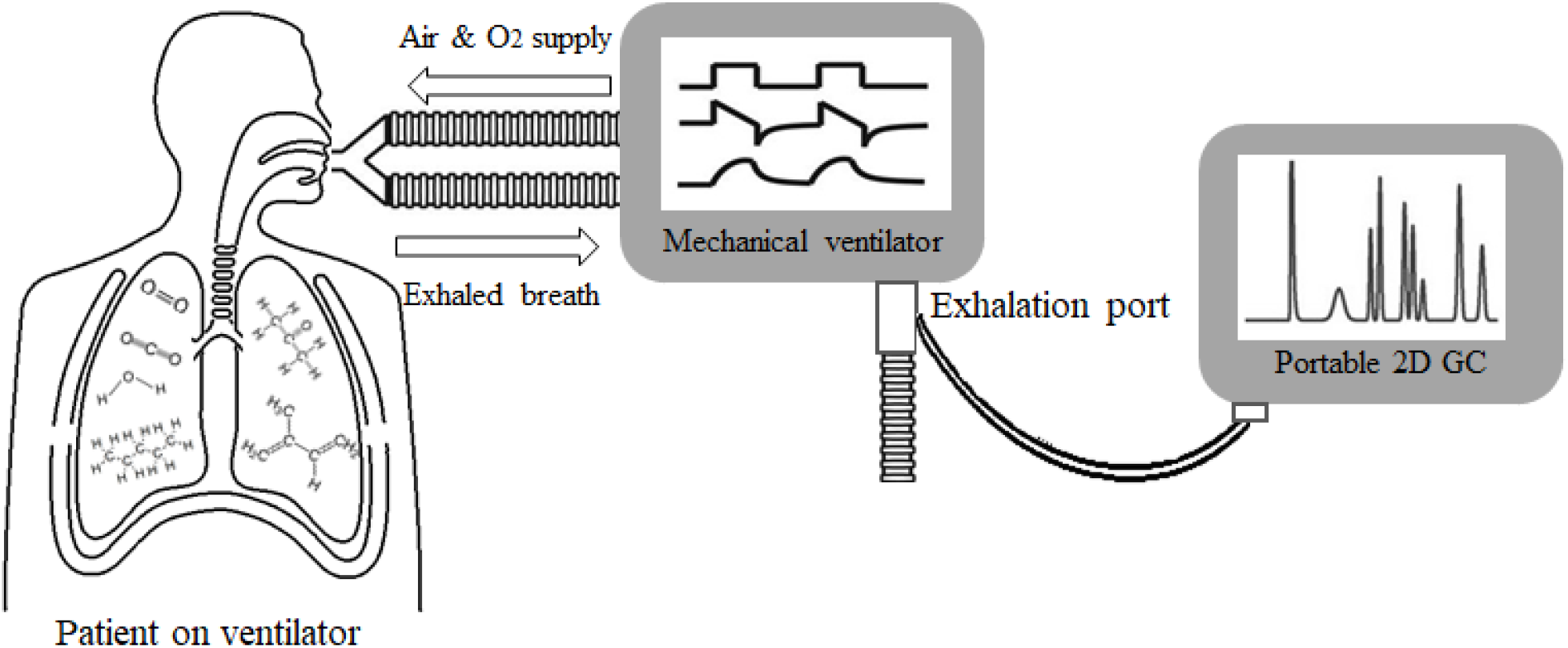
Conceptual illustration of using a portable GC device to analyze breath from a patient on a mechanical ventilator.

## 2. Methods

### 2.1. Patient selection and diagnosis

This study was approved by the University of Michigan Institutional Review Board (IRB) to consent adult patients 18 years or older undergoing mechanical ventilation for both hypoxic and non-hypoxic respiratory failure or need for mechanical ventilation for other life-support issues in various intensive care unit settings. Etiologies for the need for intubation and mechanical ventilation included ARDS, pneumonia, sepsis, pulmonary embolism, traumatic brain injury, cardiac arrest, severe chronic obstructive pulmonary disease exacerbations, and combinations of several of these reasons.

The final diagnosis of ARDS was adjudicated retrospectively by a multi-physician panel blinded to portable GC data. The adjudication was based on the Berlin Criteria^9^, which relies on a combination of medical history, chest radiography findings, and oxygenation parameters^9^. Details regarding this ARDS adjudication process have been previously reported^35^.

An adjudication was performed for the day the patient was studied using the portable GC. If a patient was studied at more than one-time point, a separate adjudication was made on those days. The adjudication of ARDS was binary (present/not present) and no attempt was made to score ARDS (if present) as mild, moderate, or severe. If patient subjects were successfully liberated from mechanical ventilations, no additional GC testing was performed.

In order to identify and populate the algorithm with breath signatures from individuals with no acute illness or injury requiring mechanical ventilation, we also recruited five laboratory members with no history of pulmonary conditions or acute illness as volunteer controls (denoted as Patient #1, 2, 3, 4, and 38 in Figures 7, S4, and S6), which serve as the baseline for the algorithm. Their breath samples were collected in Tedlar bags through a moisture filter and then immediately withdrawn into the GC device for analysis. Patient #1, 2, 3, and 4 were used in the training set whereas Patient #38 was used in the testing set.

A total of 21 ARDS patients and 27 non-ARDS control patients were recruited for 85 sets of breath chromatograms.

**IRB Statement:** This clinical research study (HUM00103401) was approved by the University of Michigan Medical School’s Institutional Review Board (a component of the University of Michigan’s Human Research Protection Program). Consent was required from patient subjects or their legally authorized representative prior to enrollment.

### 2.2. Separation and detection of breath with the portable GC device

As shown in Figure 2 patient’s breath was collected via a 2 m long polytetrafluoroethylene (PTFE) tubing (0.64 cm i.d.) connected to the exhalation port of the ventilator. The sampling rate was 70 mL/min and the sampling time was 5 minutes. The total assay time was 33 minutes, which included 5 minutes of sample collection time, 5 minutes of desorption/transfer time, 13 minutes of separation time, and 10 minutes of cleaning time.

**Figure 2.**
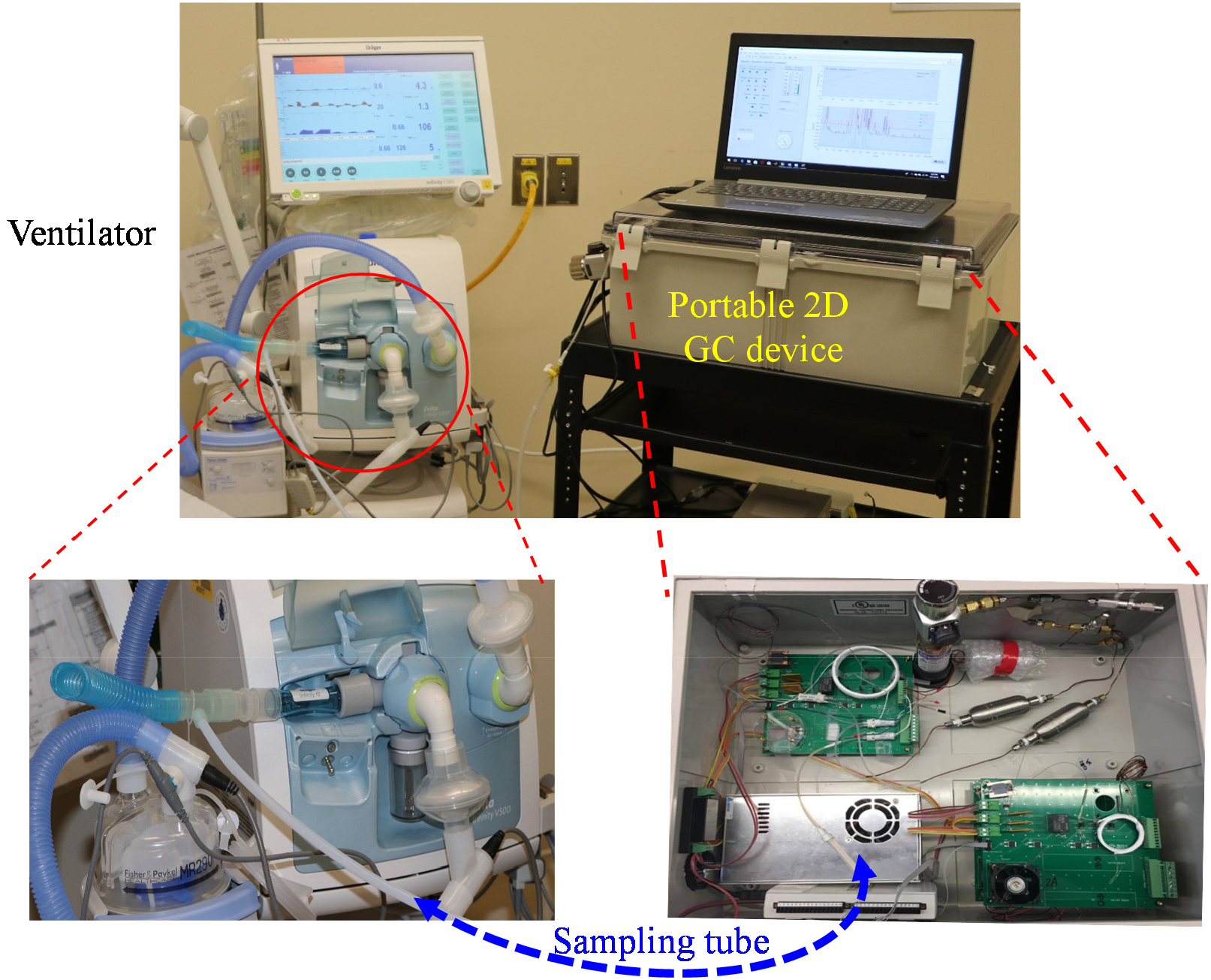
Detailed description of the experimental setup. The portable GC was connected to the output of a ventilator via a 2 m long polytetrafluoroethylene (PTFE) tubing (0.64 cm i.d.). The portable GC weighed less than 5 kg. The patient breath was drawn into and captured by the thermal desorption tube in the GC device at a flow rate of 70 mL/min for 5 minutes. The total assay time was 33 minutes, which included 5 minutes of sample collection time, 5 minutes of desorption/transfer time, 13 minutes of separation time, and 10 minutes of cleaning time. For details of GC operation, see the Supplementary Information.

In Figure 3 we give brief description of the portable 1×2-channel 2D GC and its operation. The details can be found in Section S1 in the Supplementary Information. The 2D GC device consisted of three detachable modules: sampling module, 1^st^-dimensional separation module, and 2^nd^-dimensional separation module. The 1^st^-dimensional module further consisted of a home-made micro-thermal injector (μTI), a 10 m long on-polar DB-1ms column (250 µm x 0.25 µm, Agilent J&W Scientific), and a home-made micro-photoionization detector (μPID)^36^. The 2^nd^-dimensional module consisted of two identical channels, each of which had a 3 m long polar SUPELCOWAX^®^ 10 column (250 µm x 0.25 µm, Sigma Aldrich). Note that while polar columns have been used in the 2^nd^-dimensional column in 2D GC analysis of breath^28, 29^, mid-polar columns can also be used^30^. The 1×2-channel 2D GC can be operated as a 1D GC alone when the 2^nd^-dimensional module is either disabled or detached or as comprehensive 2D GC. To increase the separation capability, in this work we chose to operate our portable GC in a comprehensive 2D GC mode, which required additional but negligible 20 seconds compared to 1D GC operation alone. In the comprehensive 2D GC mode, eluted analytes from the ^1^D column was sliced by the micro-Deans switch, loaded onto the one of the μTIs (μTI 2A or μTI 2B in Figure 3), and then injected into the corresponding ^2^D column (^2^D column 2A or ^2^D column 2B in Figure 3). The modulation time was 10 seconds so that the maximally allowed separation time on each ^2^D column was 20 seconds^34^. The ^1^D column was programmed to start from 25 °C for 2 min and was then heated to 80 °C with a ramping rate of 10 °C/min. Then the temperature was raised to 120 °C with a ramping rate of 20 °C/min and finally kept at 120 °C for 4 min. Both ^2^D columns were kept at 75 °C. In our 1×2-channel 2D GC, we used 3 flow-through μPIDs, one at the end of the ^1^D column (μPID 1 in Figure 3) and two at the end of the ^2^D columns (μPID 2A and μPID 2B in Figure 3). The use of a detector at the end of ^1^D column allows us to monitor the elution of the analytes from the ^1^D column to produce the 1D chromatogram (if the GC device is operated as 1D GC alone) or to avoid potential under-sampling that may be caused by the 10 s modulation time (if the GC device is operated as comprehensive 2D GC) (see the detailed discussion in Ref. 34).

**Figure 3.**
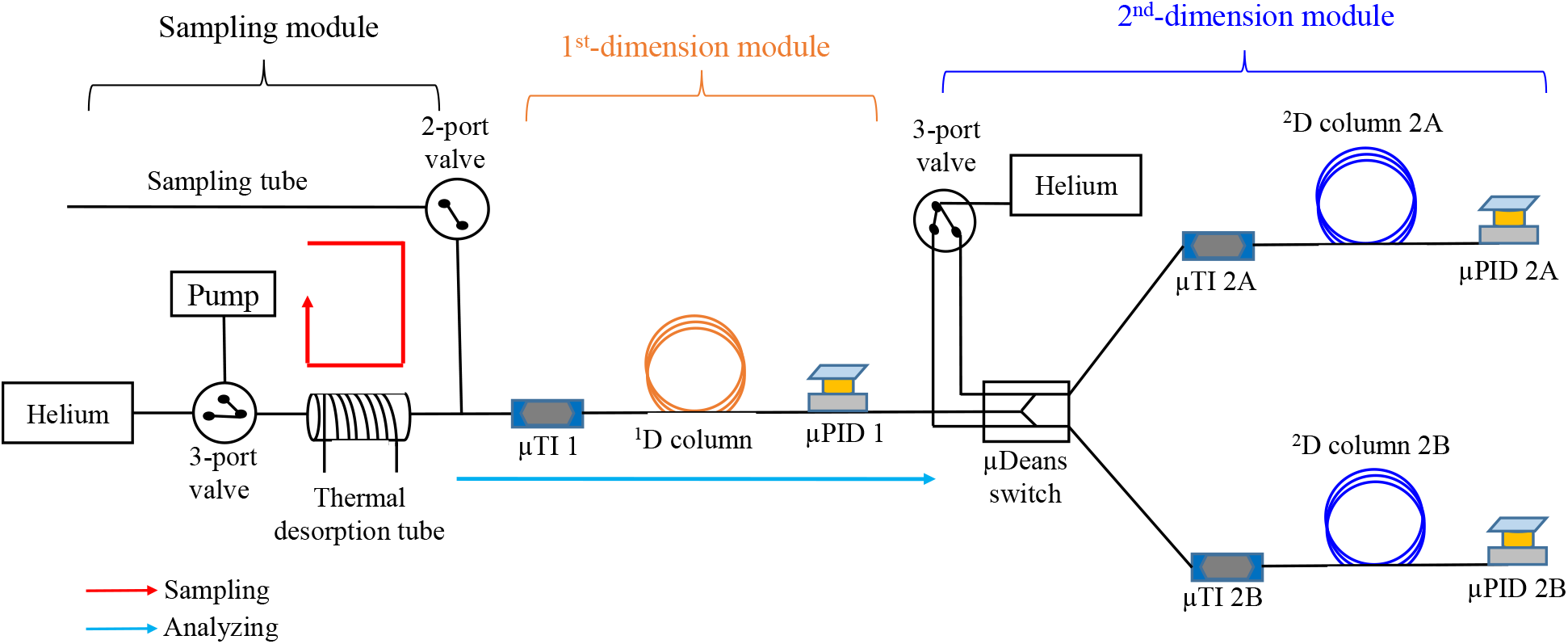
Layout of the portable 1×2-channel 2D GC device. It consisted of three detachable modules: sampling module, 1^st^-dimensional separation module, and 2^nd^-dimensional separation module. The 1^st^-dimensional module consisted of a micro-thermal injector (μTI), a non-polar column, and a micro-photoionization detector (μPID). The 2^nd^-dimensional module had two identical channels, each of which consisted of a μTI, a polar column, and a μPID. During operation, breath was collected via the sampling tube and captured by the thermal desorption tube. Then the analytes were transferred to μTI 1 and injected into the ^1^D column. The elution from the ^1^D column was detected by μPID 1. If 2^nd^-dimensional separation was needed, the 2^nd^-dimensional module was attached to the outlet of the ^1^D column and the elution from the ^1^D column was sliced and sent alternately to one of the two ^1^D columns via a micro-Deans switch. The portable GC can operate as 1D GC alone when the 2^nd^-dimensional module is disabled or detached, or as comprehensive 2D GC. Comprehensive 2D GC operation required additional 20 seconds compared to 1D GC operation alone, which was negligible considering the overall assay time of ∼30 minutes. For details of GC operation and 2D GC chromatogram construction, see the Supplementary Information.

Details of 2D GC chromatogram construction based on the signal obtained by the 3 detectors and the algorithm for the subsequent analysis are described in Sections S2 and S3, respectively.

## 3. Results

### 3.1. Chromatograms for ARDS and non-ARDS patients

Figure 4 shows the representative 1D and 2D chromatograms for an ARDS and a non-ARDS control. It can be seen from Figures 4 and S1 that 2D GC provides additional separation capability as compared to the 1D GC (3-10 times higher in terms of peak capacity according to the analysis in Section S4 in the Supplementary Information). Section S4 further compares the separation capability between our 2D GC and other technologies such as IMS and SIFT-MS. Figure 5 shows that a total 97 peaks were found collectively in the 85 2D chromatograms from the patients under study, among which nearly 70% of the peaks are co-eluted or partially coeluted in the ^1^D column. Note that not all 97 peaks appear in a 2D chromatogram for a particular patient, as some peaks are below the detection limit of our μPIDs. Recent studies using commercial bench-top comprehensive 2D GC coupled with time-of-flight mass spectrometry identified approximately 2000 VOC peaks in a 2D chromatogram^30^. Although our portable 2D GC does not generate as many peaks as the high-end bench-top GC-MS, it can still distinguish ARDS and non-ARDS as shown later. Also, among all the recruited and adjudicated patients, we were able to monitor 9 ARDS patients and 9 non-ARDS patients for multiple days. Figure 6 shows, as an example, the 2D chromatograms for an ARDS patient tested over 3 days, from which we can see clearly that there are both qualitative and quantitative changes in breath VOCs.

**Figure 4.**
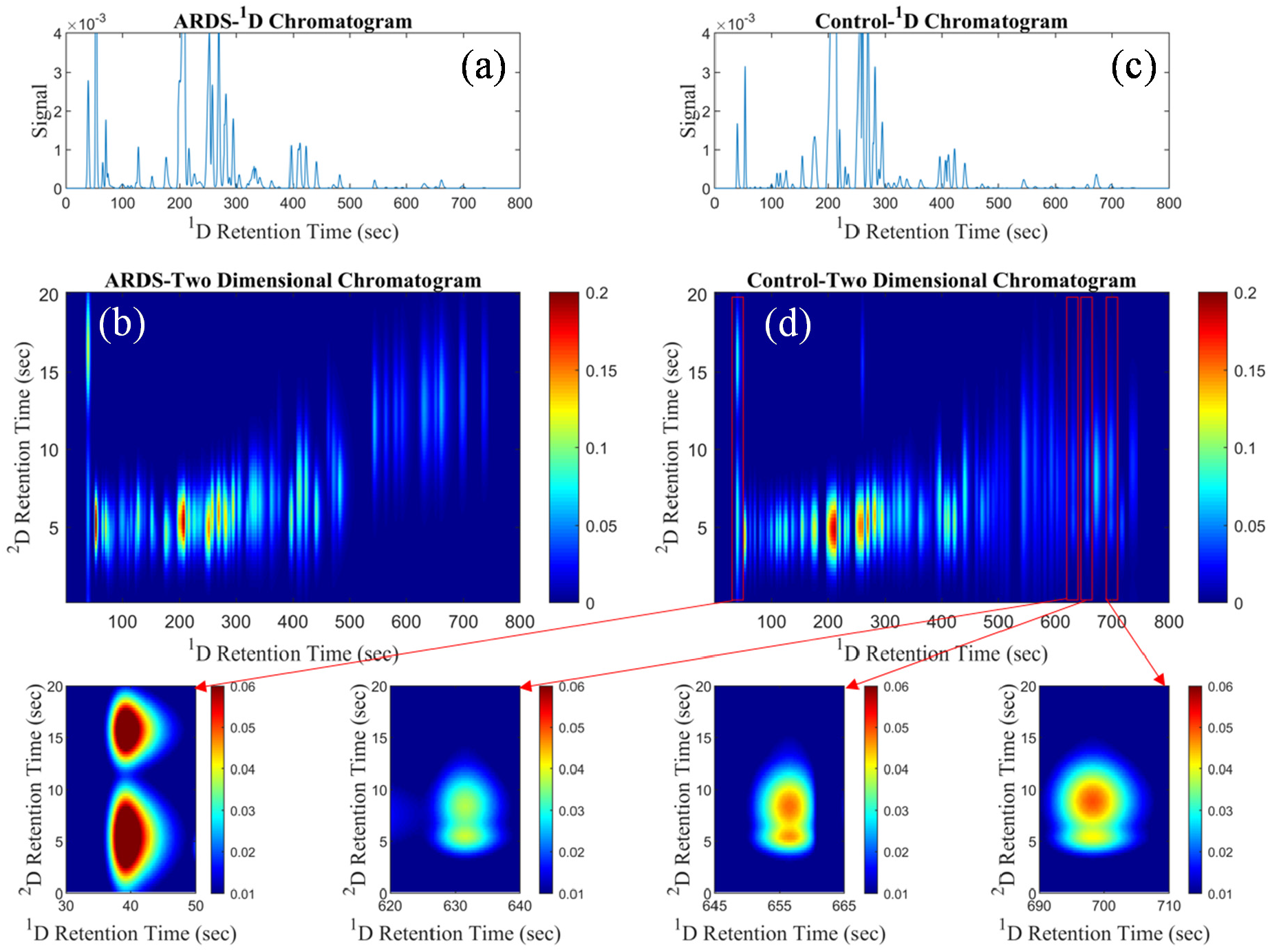
(a) and (b) Representative 1D chromatogram and 2D chromatogram for an ARDS patient, respectively. (c) and (d) Representative 1D chromatogram and 2D chromatogram for a non-ARDS (control) patient, respectively. In the zoomed-in 2D chromatogram for the control patient, four co-eluted ^1^D peaks are separated into eight peaks in the 2D chromatogram. Other zoomed-in portions of (b) and (d) can be found in Figure S1 in the Supplementary Information.

**Figure 5.**
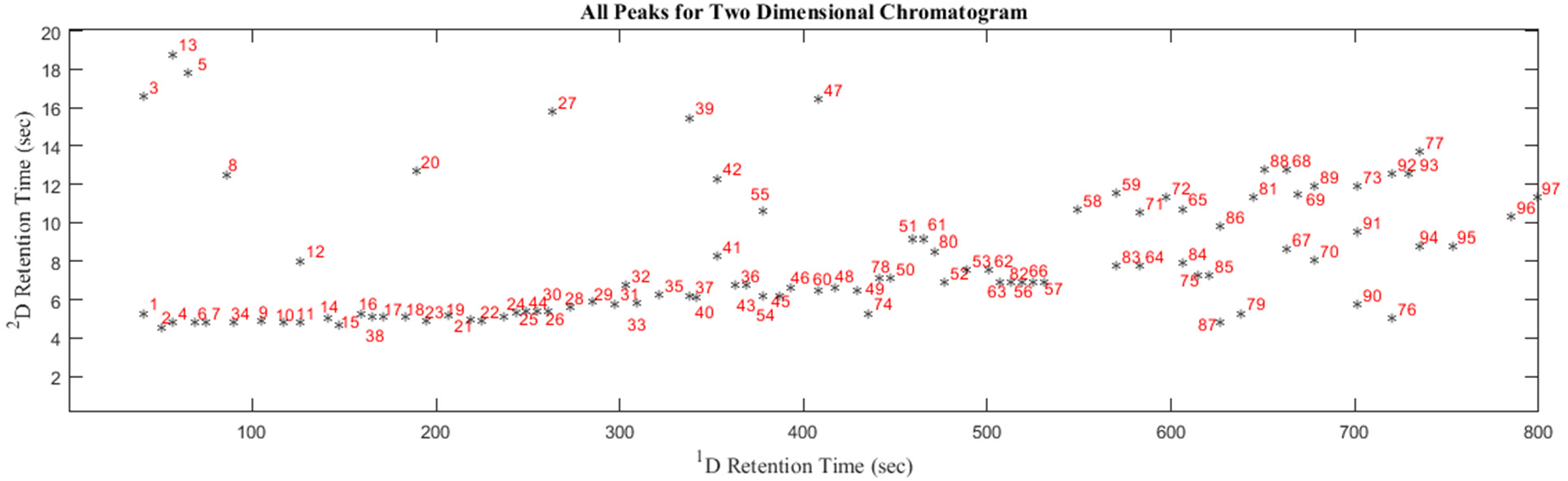
All 97 peaks found collectively in 85 breath samples from 48 patients plotted in a 2D chromatogram, among which 18 pairs (36 peaks) are co-eluted and approximately another 30 peaks are partially co-eluted (with doublets or triplets and separation of adjacent peaks is less than 2σ) from the ^1^D column. Each dot represents the center of a peak in the contour plot (see, for example, Figure 4, for a peak contour plot). Note that not all 97 peaks appear in a 2D chromatogram for a particular patient.

**Figure 6.**
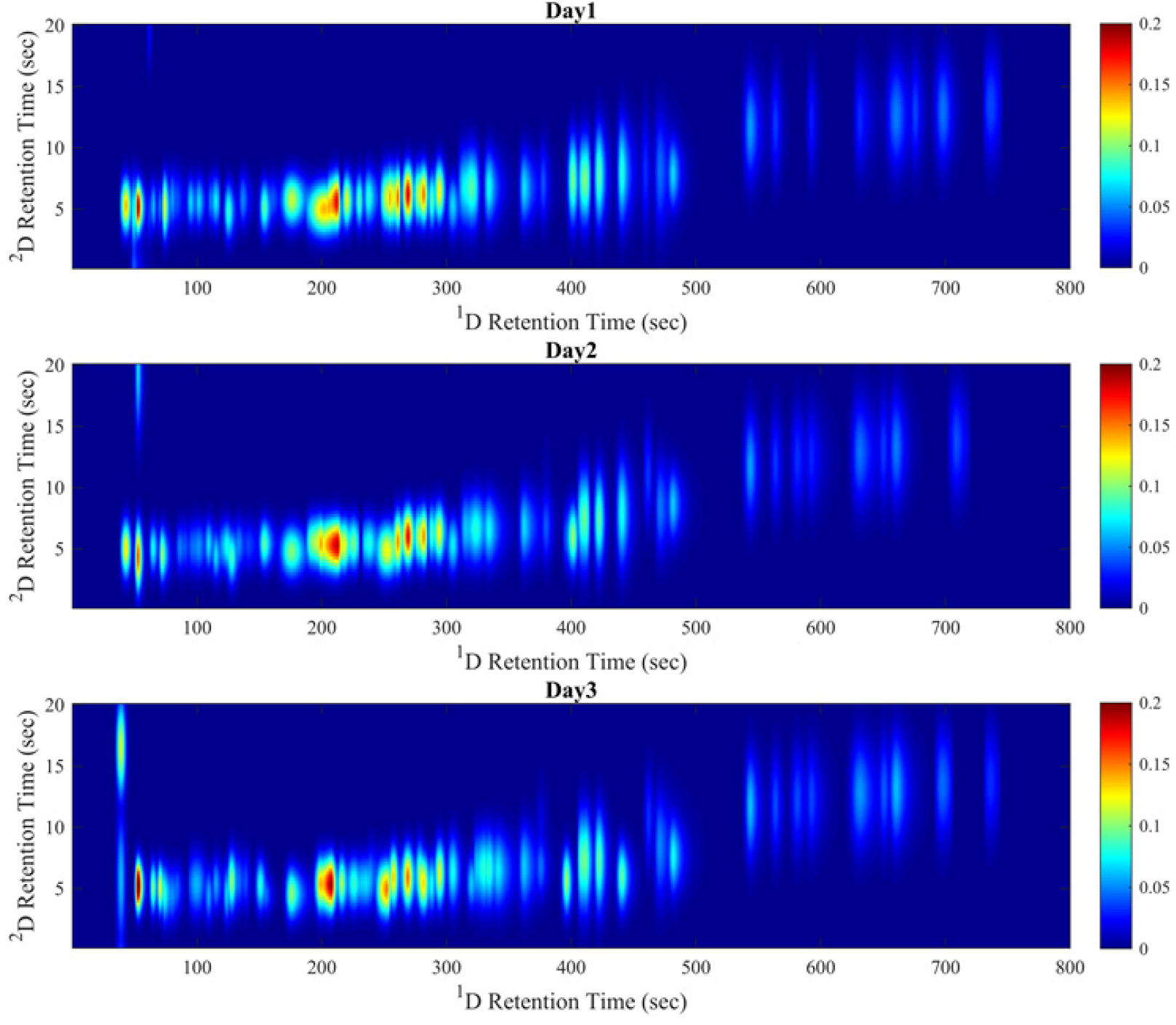
Evolution of the 2D chromatogram of an ARDS patient (Patient #11) during 3 days of monitoring.

### 3.2. Patient classification based on 2D chromatograms

Among all 97 peaks, not all of them appear to be relevant to ARDS. For example, some of the peaks may be from indoor air background, normal metabolic activities, or other conditions that a patient may have. Those irrelevant peaks interfere in the correct classification of ARDS and non-ARDS groups. It is therefore critical to determine which subset of the peaks is most responsible for the differences observed between ARDS and non-ARDS groups. To select the optimal subset of peaks, 28 patients (11 ARDS, 17 control, and a total of 43 tests) were used as the training set, whereas the remaining 20 patients (10 ARDS, 10 controls, and a total of 42 tests) were used as the testing set.

An algorithm based on machine learning, principal component analysis (PCA), and linear discriminant analysis (LDA) was developed to first select the optimal subset of peaks using the training set. Then with the PCA coefficient acquired from the training set, the PCA scores for the testing set of patients can be calculated. The details of the algorithm are given in Section S3 in the Supplementary Information. With this algorithm, we selected 9-peak subset as the final optimal peak subset, which yields the best classification accuracy (93.0%) and the maximum boundary distance. The final PCA scores for all recruited patients are shown in Figure 7. The final PCA model achieved an overall accuracy of 87.1% with 94.1% positive predictive value and 82.4% negative predictive value. The corresponding specificity, sensitivity, positive predictive value (PPV), and negative predictive value (NPV) are presented in Table 2. The corresponding Q-residuals for all recruited patients are shown in Figure S5. Separate PCA scores for the training and testing sets the corresponding statistics (specificity, sensitivity, PPV and NPV) are presented in Figure S6 and Table S2, respectively.

**Figure 7.**
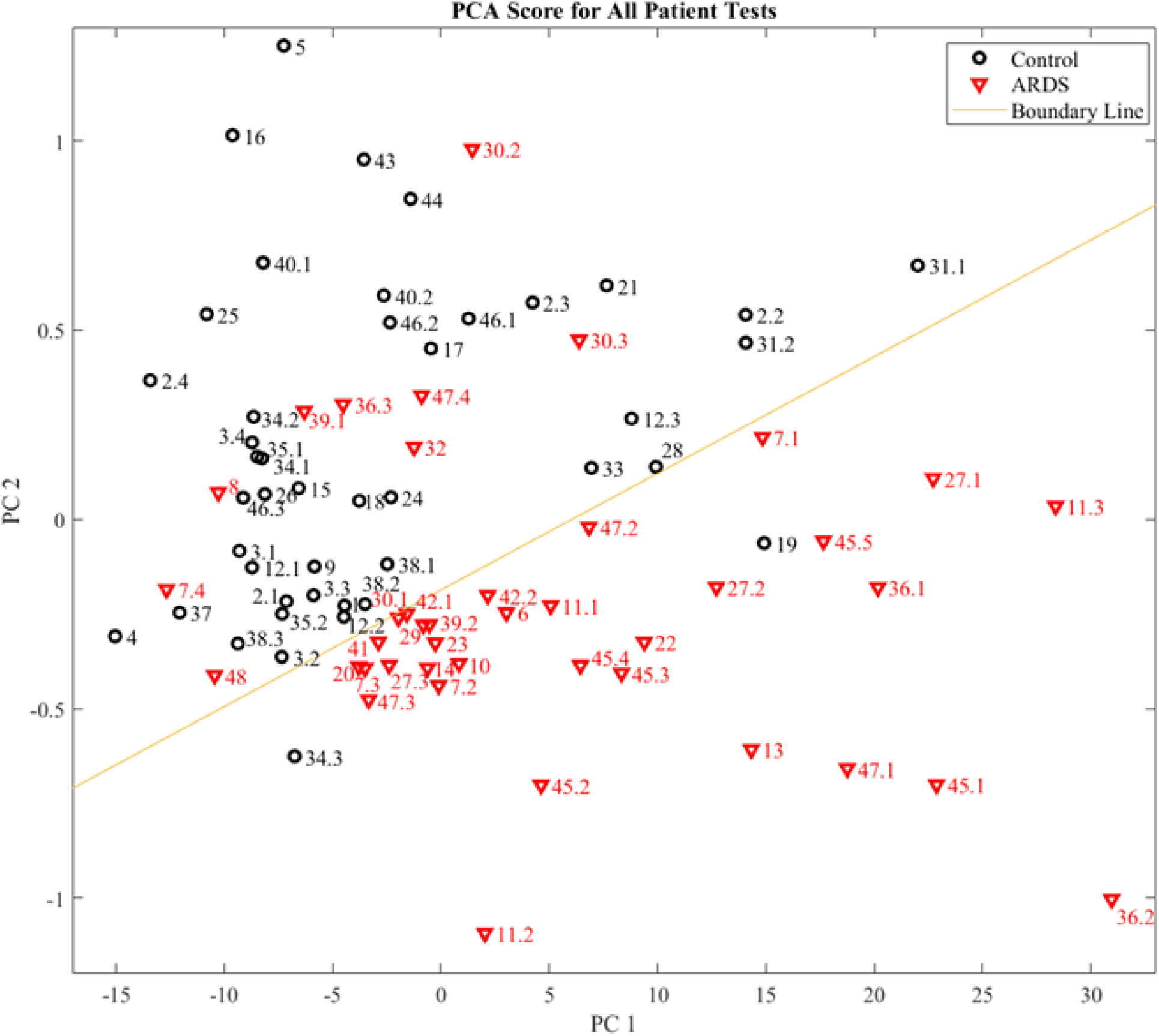
PCA plot of all recruited patients. X-axis (PC 1) is the 1^st^ principal component and Y-axis (PC 2) is the 2^nd^ principal component. The red and black symbols denote respectively the ARDS and non-ARDS patients adjudicated by physicians using the Berlin criteria. The patient numbers are given by the symbol. For example, “11.1” and “11.3” denote Patient #11, Day 1 and Day 3 results, respectively. The bottom/top zone below/above the boundary line represents respectively the ARDS/non-ARDS region using the breath analysis method. The corresponding Q-residuals for this PCA model are shown in Figure S7.

**Table 2.** Statistics of breath analysis for ARDS.

|  | **ARDS** | **Non-ARDS** | **Total** |
| --- | --- | --- | --- |
| **Positive (our results)** | 32 | 2 | 34 |
| **Negative (our results)** | 9 | 42 | 51 |
| **Column total** | 41 | 44 | 85 |
| **Specificity** | 1-2/44 =95.5 % |  |  |
| **Sensitivity** | 1-9/41=78.0 % |  |  |
| **Positive predictive value** | 32/34 =94.1 % |  |  |
| **Negative predictive value** | 42/51= 82.4 % |  |  |
| **Total accuracy** | 74/85= 87.1% |  |  |

### 3.3. Time series measurement of ARDS patients

One of important advantages of breath analysis is the potential to non-invasively monitor the development of ARDS, the severity of ARDS (if present), and the resolution of ARDS. This would allow the technology to map the trajectory of the disease and potentially guide therapy and decision making. Among the 9 ARDS patients and 9 non-ARDS patients whom we monitored on multiple days, some ARDS patients were noted to clinically progress to a non-ARDS status and vice versa, as determined by both 2D GC and clinical adjudication. Our results demonstrated that breath analysis may be able to predict the ARDS trajectory 12-48 hours in advance. Below we show some examples, whose trajectories on the PCA plot are shown in Figure 8.

**Figure 8.**
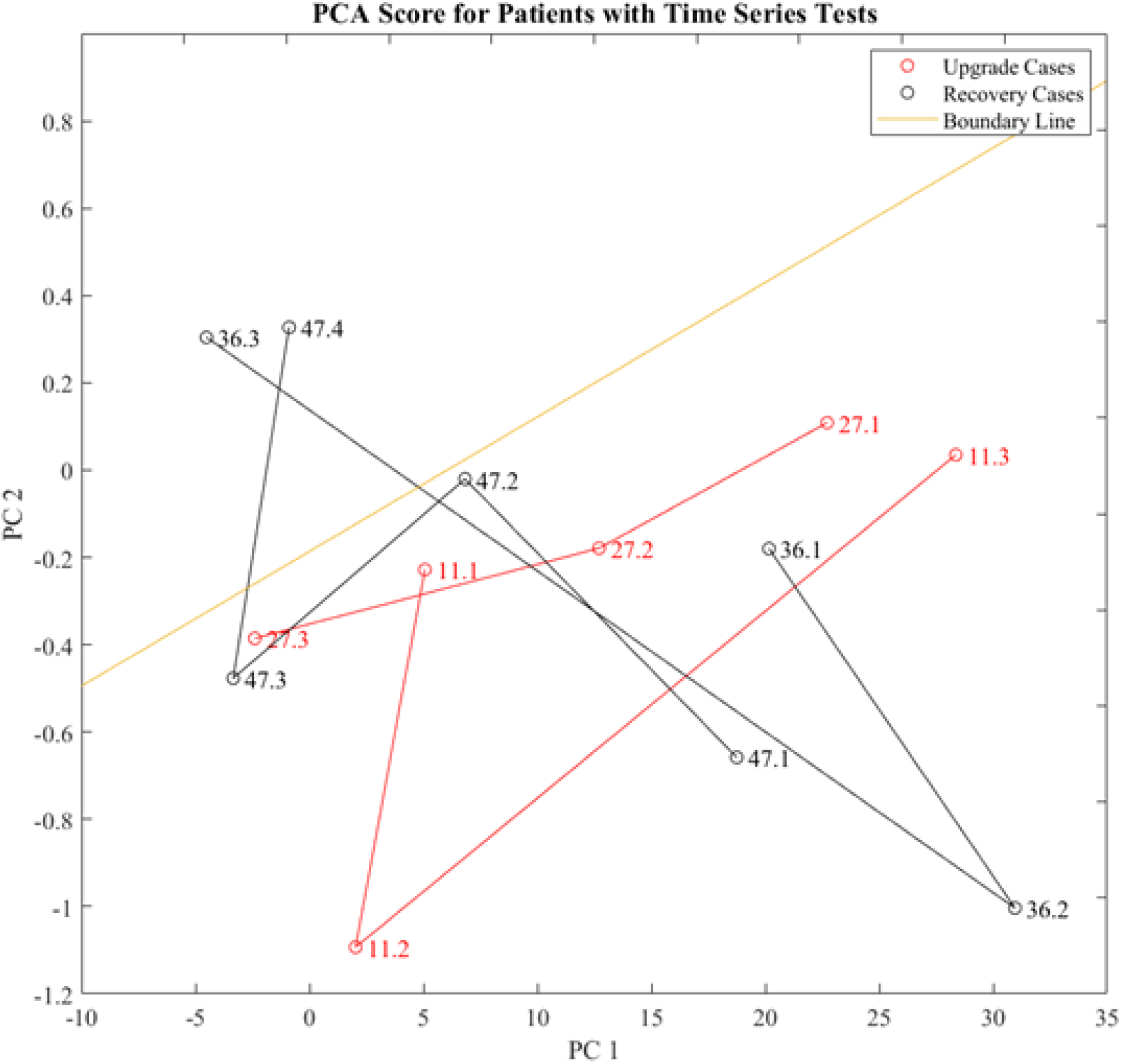
The trajectory on PCA plot for patient #11, #27, #36, and #47. #11 and #27 are the upgrade case (initially listed as potential ARDS on the first day) and #36 and #47 are recovery cases (extubated and discharged from ICU 24-48 hours after the last test). The bottom/top zone below/above the boundary line represents respectively the ARDS/non-ARDS region using the breath analysis method.

#### (1) Upgrade cases

Patient #11 was a potential and undetermined ARDS patient (meaning that the clinician suspected that the patient might develop ARDS, but was not certain at the time of diagnosis. The patient was placed on close monitoring) on the 1^st^ test day and then upgraded to ARDS on the next day. The breath test suggested ARDS from the 1^st^ test day to the 3^rd^ test day (#11.1, #11.2, and #11.3).

Patient #27 was a potential and undetermined ARDS patient on the 1^st^ test day and then upgraded to ARDS on the next day. The breath test suggested ARDS from the 1^st^ test day to the 3^rd^ test day (#27.1, #27.2, and #27.3).

#### (2) Recovery cases

Patient #36 was sampled for 3 days. On the 3^rd^ day the patient was still listed as ARDS patients based on the Berlin Criteria and then got extubated (liberated from mechanical ventilation) and discharged from ICU on the 5^th^ day. The breath tests for the first 2 days suggested ARDS (#36.1 and #36.2). The breath test for the 3^rd^ day demonstrated a non-ARDS pattern (#36.3).

Patient #47 was sampled for 4 days and was liberated form mechanical ventilation and discharged on the 6^th^ day. Based on the Berlin Criteria this patient had ARDS for all first 4 days. The breath tests for the first 3 days breath tests show an ARDS pattern (#47.1, # 47.2, and #47.3) and the 4^th^ day breath test shows as non-ARDS pattern (#47.4).

Note: If the breath test results do not match the clinical adjudication, we consider the test as “mis-classification” when calculating the overall classification accuracy, even for the cases like #36.3 and #47.4 that suggest that the breath analysis was able to predict the trajectory of ARDS (i.e., earlier diagnosis).

With further evidence on the following days, the two potential and undetermined ARDS cases mentioned above (#11.1 and #27.1) were finally determined as ARDS based on the Berlin Criteria.

The trajectories of the entire 18 patients and their medical histories can be found in Figure S7 and Section S5.

## 4. Discussion

To our knowledge, the portable 2D GC device described here is the first of its kind that can be used in POC to continuously monitor patient breath. Using this portable GC device, along with algorithms developed in this work, we are able to distinguish ARDS and non-ARDS with high accuracy, compared to clinical expert adjudication. As a dynamic syndrome with multiple etiologies, the real-time diagnosis of ARDS is challenging. There are currently no technologies allowing its real-time diagnosis or tracking. The only widely available tool in use in assisting in ARDS diagnostics is LIPS. However, LIPS was designed as a screening tool that incorporates a series of risk factors and risk modifiers to predict whether ARDS will occur at a future point. While a small subset of data using the 2D GC technology indicates the potential to predict onset or resolution, much more testing will be needed. An interesting possibility would be to utilize LIPS in conjunction with the technology to improve screening^17, 19^.

It should be noted that based on the results obtained through the current work the 9-peak subset (Peak #2, 34, 38, 44, 62, 66, 72, 79, and 81 in Figure 5) that was selected for ARDS detection can be almost separated using the ^1^D column in our current 2D GC device (except Peak #34 that is nearly co-eluted with Peak #8. See Figure S3). Therefore, it seems that if we only need to distinguish between ARDS and non-ARDS, then our portable GC can simply be operated in a 1D GC mode alone. However, we believe that 2D GC operation is still a preferred choice, since the potential co-elution of Peak #34 and #8 may affect the ARDS detection. More significantly, 2D GC operation is critical to sub-typing ARDS and analyzing complications. For example, the peaks (#3, 5, 13, and 27) in Figure 5 do not belong to the ARDS-relevant 9-peak subset, but they have different concentrations between the ARDS patient and the healthy control (see Figures 4(b), 4(d), and S1), suggesting that the ARDS patient in Figure 4(b) might have other health conditions besides ARDS. In addition, the peaks (#3, 5, and 13) in Figure 6, which are not part of the 9-peak subset, indicate other health conditions (besides ARDS) of the same patient might change during medical treatment of ARDS. For future applications (in detection of ARDS and ARDS with complications, and in detection of other diseases such as asthma and pneumonia), it is still preferred to continue to use 2D GC to separate as many peaks as possible, which makes the device much more flexible for various diseases and medical conditions rather than being dedicated to monitoring of ARDS alone.

This study has a number of important limitations. First and foremost is that the histopathologic examination of lung tissue for changes consistent with diffuse alveolar damage (DAD) was not used to make the diagnosis of ARDS. Even in those patients dying of respiratory failure, autopsies were not obtained. While DAD on histopathology is the pathologic gold standard, obtaining serial lung biopsies for diagnosis is not feasible or a clinical standard of care. Although the clinical consensus for the diagnosis of ARDS is the Berlin criteria, Kao and others have demonstrated that of patients clinically diagnosed as having ARDS using the Berlin criteria, less than 60% have DAD on lung histopathology when lung biopsies can be obtained^11^. In the absence of tissue biopsy, we employed the best available instrument (multi-physician adjudication) for identifying ARDS^35^.

While this is a limitation to our study, it is also a limitation to any clinical research or clinical trial in the field of ARDS further underscoring the need for new diagnostics. Despite this, we noted the VOC patterns that where clearly distinguishable from patterns seen in subjects who were mechanically ventilated for non-hypoxic respiratory failure such as sepsis (without ARDS) and cardiac arrest, as well as those intubated for hypoxic respiratory failure in whom after intubation and mechanical ventilation the patient’s PaO_2_/FiO_2_ was clearly not indicative of ARDS (COPD exacerbation, pulmonary embolism, and unilateral pneumonia). In cases of divergence in clinical scoring and breath analysis result, differences could be due to miss-diagnosis of ARDS by clinical scoring, the ability of breath analysis to detect earlier onset or resolution of ARDS than clinical adjudication, mixed lung and systemic pathologies existing in the same patients, and others.

Finally, we note that in the current study only 48 patients (and 85 sets of breath samples) were used for the proof of the concept. A much larger groups of patients need to be recruited to further validate our method. Additionally, the selected 9 peaks as well as other peaks that may be relevant to lung and systemic pathologies should all be identified in order to understand the underlying physiological conditions of the patients and how those VOCs are involved in the ARDS processes.

## 5. Conclusions

We have developed an automated portable 2D GC device and a corresponding algorithm for breath analysis that begins to enable distinguishing the condition of ARDS from non-ARDS. Particularly, the 94.1% positive predicative value suggests that our breath analysis method can accurately diagnose ARDS, which is critical to its treatment. In several subjects studied, the technology was found to indicate the presence of ARDS prior to the development of traditional indicators used for ARDS diagnosis opening up the potential for earlier interventions. The non-invasive nature of breath analysis may also allow for continuous monitoring of ARDS trajectory as evidenced by several subjects who demonstrated changing breathomic patterns from ARDS to non-ARDS status prior to changes in traditional indicators.

The potential to leverage exhaled breath for the identification of breathomic patterns used for early diagnosis, disease trajectory tracking, and outcome prediction monitoring of ARDS would have significant impact on changing practice and improving patient outcomes. The device is envisioned to be used for ARDS patients in emergency departments, operating rooms, and intensive care units. Additionally, our device holds the potential to dramatically improve the molecular characterization of ARDS and its competing diagnoses. This clinical ambiguity of the diagnosis of ARDS compared with histopathology impairs the field’s ability to develop and study targeted, disease-specific therapies. Exhaled breath VOC analysis could significantly enhance our molecular phenotyping of patients with hypoxic respiratory failure, crystallizing our diagnoses, and dramatically improving our ability to tailor treatments to and enrich clinical trials with patients with true ARDS pathophysiology.

## Author contributions

X. F. and K. R. W. take responsibility for the content of the manuscript, including the data and analysis. Study concept and design: K. R. W., X. F., M. Z., R. S., and H. Z.; acquisition, M. Z., R. S., H. Z., J. L., S. W., E. B., J. M., A. P.; analysis or interpretation of data: M. Z., R. S., M. S., R. P. D., K. R. W., and X. F.; administrative, technical, or material support: E. B., J. M., A. P., R. H., and L. N.; obtained funding: X. F., K. R. W., and R. P. D.; manuscript writing: M. Z., R. S., M. S., R. P. D., P. P., K. R. W., and X. F.; agreeing to be accountable for all aspects of the work in ensuring that questions related to the accuracy or integrity of any part of the work are appropriately investigated and resolved: all authors.

## Acknowledgments

The authors acknowledge the financial support from National Institutes of Health (1-R21-HL-139156-01), NIH Center for Accelerated Innovations at Cleveland Clinic (Program Prime Award Number: 1UH54HL119810-05; Project Number: NCAI-17-7-APP-UMICH-Fan), the Michigan Translation and Commercialization (MTRAC) for Life Sciences Hub, and the Michigan Center for Integrative Research in Critical Care. They also thank the support from Flux HPC Cluster provided by the University of Michigan Office of Research and Advanced Research Computing – Technology Services (ARC-TS).

## Supplementary information

2D GC description and operation. 2D GC chromatogram construction. Description of algorithm. Patient medical history.

