## Supplementary Information for "Rapid breath analysis for acute respiratory distress syndrome diagnostics using a portable 2-dimensional gas chromatography device"

### Section S1: Portable 2D GC Description and Operation

#### *S1.1. Materials*

DB-1ms Agilent J&W, nonpolar column (length 10 m, i.d. 250 μm, film thickness 0.25 μm) was purchased form Agilent Technologies (P/N: 122-0162, Agilent Technologies). SUPELCOWAX® 10 polar column (length 3 m, i.d. 250 μm, film thickness 0.25 μm) was purchased from Sigma Aldrich (P/N: 24077, Sigma-Aldrich). Copper tube (length 10 cm, i.d. 1 mm, o.d. 1.5 mm) was purchased from Swagelok and glass wool was purchased from Sigma Aldrich. Teflon tape was purchased from Grainger (Ann Arbor, MI). Shrink tube was purchased form Digi-Key Electronics. Other materials are the same as those described in Ref. 1.

#### *S1.2. Design, fabrication, and characterization of components*

Various microfabricated components were used in the present portable 1 x 2 channel 2D GC device, including thermal desorption tube, micro-fabricated thermal injector (μTI), micro-Deans switch (μDS), and micro-photoionization detector (μPID). All of these components were fabricated and characterized in-house. The details of μTI, μDS, and μPID can be found in Ref. 1. The detail of the thermal desorption tube is given as follows.

The thermal desorption tube was made of a 5 cm long copper tube having an inner diameter of 1 mm. Both Carbopack™ X and B granules, 10 mg each, were loaded into the hollow cylindrical copper tube using a diaphragm pump. Glass wool was used to separate the Carbopack™ X and B, as well as to seal the copper tube from both ends. Swagelok fittings were used to connect a stainless steel tube of i.d. 250 μm at both the ends of the copper tube. For temperature ramping, the nickel wire was wrapped around the entire length of the copper tube. The nickel wire was insulated from the copper tube using a Kapton tape. A type K thermocouple was attached to the copper tube using a Kapton tape to monitor the temperature in real time. Finally, the thermal desorption tube was preconditioned at 300 °C for 12 h under helium flow.

#### *S1.3. Device assembly*

The portable 1 x 2 channel 2D GC device is similar to the 1 x 4 channel 2D GC device described in Ref. 1. As illustrated in Figure 3 in the main text, the 2D GC consisted of a sampling module, a 1$^{st}$-dimensional separation module, and a 2$^{nd}$-dimensional separation module. The sampling module consisted of a sampling tube, a thermal desorption tube loaded with Carbopack™

X and B, valves, and a pump. The 1st-dimensional module consisted of a μTI loaded with Carbopack™ X and B, a 10 m long Agilent J&W DB-1ms, and a μPID. The 2nd-dimensional module had two identical channels. Each channel consisted of a μTI, a 3 m long SUPELCOWAX® 10 column, and a μPID. The eluent from the $^{1}$D column was transferred to one of the $^{2}$D columns via a μDS. All the modules and components were connected via tubings, universal connectors, and Y-connectors. The entire device was housed in a customized plastic case (see Figure 2 in the main text) and had a total weight less than 5 kg, including the weight of the He gas cartridge (231 g). LabVIEW™ based codes were developed in-house for user interface, and device control and automation.

The portable GC can be operated in 1D GC alone (in which case the 2nd-dimensional module was disabled or detached) or comprehensive 2D GC. Operation as 1D GC is straightforward. Operation as comprehensive 2D GC is described in Section S1.4 below.

During the measurement, the device was secured on a rolling cart (see Figure 2) and placed outside the ventilated patient's room. A 2 m long polytetrafluoroethylene (PTFE) tubing (0.64 cm i.d.) was used to connect the output of the ventilator to the GC device for breath sampling.

***S1.4. Operation of the portable 1x2-channel 2D GC***

The operation procedures and parameters of the portable GC in the comprehensive 2D GC mode are described as follows.

(1) Sampling: The exhaled breath of the patient was drawn by the diaphragm pump through the 2-port valve and adsorbed by the thermal desorption tube at a flow rate of 70 mL/min for 5 min.

(2) Desorption and injection: The 2-port valve was closed and the helium gas was flowed through the 3-port valve at a flow rate of 2 mL/min. Meanwhile, the thermal desorption tube was heated to 300 °C for 5 min to transfer the analytes onto the μTI 1. After 5 min, μTI 1 was heated to 270 °C in 0.6 s and then kept at 250 °C for 30 s for complete thermal desorption and injection of the analytes into the $^{1}$D column.

(3) Separation: The analytes underwent separation through the 10 m long $^{1}$D column and then detected by μPID 1. During the separation, the column was kept at 25 °C for 2 min, then first ramped at a rate of 10 °C min$^{-1}$ to 80 °C, next ramped at a rate of 20 °C min$^{-1}$ to 120 °C, and kept at 120 °C for 4 min. The flow rate was 2 mL/min for the $^{1}$D column.

We used a modulation period of 10 s to inject the eluent from the $^{1}$D column into the $^{2}$D columns. The first 10 s long slice of the eluent from the $^{1}$D column was routed to and trapped by μTI 2A, which were kept at room temperature (25 °C). The μTI 2A was then heated to 270 °C in 0.6 s and then kept at 250 °C for 5 s to inject the trapped analytes to Column 2A. In the meantime, the second 10 s long slice of the eluent from the $^{1}$D column was routed to and trapped by μTI 2B, which was subsequently injected into Column 2B. The same operation was repeated between Columns 2A and 2B alternatively throughout the analysis. The analyte underwent $^{2}$D separation through one of $^{2}$D columns (kept isothermally at 75 °C during entire operation) and then detected by μPID 2. During the entire operation, the helium flow was 3 mL/min for each of $^{2}$D columns.

(4) Cleaning: After analysis, the outlet of the μTI 1 was disconnected from the inlet of the $^{1}$D column so that it was open to the ambient air. Then the thermal desorption tube was heated to 300 ºC for 5 min followed by heating μTI 1 to 270 °C in 0.6 s and then kept at 250 °C for 30 s at a helium flow rate of 25 mL/min. This process was repeated twice in order to completely remove the residual analytes (if any) trapped in the thermal desorption tube and μTI 1. Note that cleaning of μTI 2A and 2B was not needed.

The total assay time was 33 minutes, which included 5 minutes of sample collection, 5 minutes of desorption/transfer, 13 minutes of separation, and 10 minutes of cleaning.

It should be noted that multiple μPIDs were used to measure the analytes eluted from $^{1}$D column and $^{2}$D columns. The responsivity of those μPIDs may be different due to variations in aging and amplification, etc. During the experiment, μPID 2A and 2B were calibrated against μPID 1 using toluene (50 ppb), as discussed in detail in Ref. 1. This calibration was carried out approximately every 300 hours of operation.

Operation of the portable GC as 1D GC is similar to the steps in (1)-(4) above, except that the inlet of the μDS is detached from the outlet of the μPID 1 or the 2$^{nd}$-dimensional module is powered off.

### Section S2: Two-dimensional gas chromatogram construction

Once the analysis is completed, two-dimensional gas chromatogram was generated from all three channels' (1D, 2A, 2B) PID signals.

First, the signal from each channel was preprocessed for baseline correction and peak detection. Second, all the $^{2}$D peaks was traced back to their correspondence $^{1}$D peak based on the $^{1}$D period they were sampled from, so that both $^{1}$D and $^{2}$D retention time and peak shapes could be found. Third, the $^{1}$D chromatogram was aligned to the reference chromatogram to fix the $^{1}$D retention time drift. Finally, the two-dimensional gas chromatogram for each peak was generated by multiplying its $^{1}$D peak shape to its $^{2}$D peak shape. Adding all individual peak's two-dimensional gas chromatogram yielded the completed two-dimensional gas chromatogram. All algorithms were implemented in the Matlab™ programming environment with a user-friendly graphical interface.

(1) Baseline correction: The baseline of gas chromatograms drift slowly due to column bleeding at high temperature, flow fluctuation and detector performance. This baseline drift can negatively affect the analytical results quantitatively hence should be corrected before performing further data analysis[2]. Here we use adaptive iterative reweighted Penalized Least Squares (airPLS) algorithms, developed by Zhang et al.[3], which iteratively changing weights of sum squares errors (SSE) between the original signals and fitted baseline until the termination criteria is met. This method requires no user intervention and has been applied to chromatograms, NMR and Raman spectra.

(2) Peak detection: After the baseline correction, peak detection is applied to both $^{1}$D and $^{2}$D chromatograms. The peak apexes are found via the method developed by Morris et al.[4]. In this method, the signal is first denoised via wavelet regression using the undecimated discrete wavelet transform (UDWT), then searched for all local maxima and the associated peak endpoints. At last, the peaks that do not meet the peak height and FWHM criteria are eliminated [4-8]. Once the peak apexes are found (including single and co-eluted peaks), the peak shapes are fitted by the exponentially modified Gaussian (EMG) model. This peak fitting method has been described previously[1].

(3) Peak assignment: Within each modulation, the $^{1}$D peak will be injected into either a 2A or 2B subsystem. Each $^{2}$D peak is assigned to one or multiple $^{1}$D peak IDs, depending on the total

peaks within each modulation. For each individual peak, multiplying its $^{1}$D peak shape by its normalized $^{2}$D peak shape yields a two-dimensional chromatogram for this individual peak.

(4) Peak alignment: Gas chromatograms may contain distortions of the retention time due to column aging, changes in temperature, or other unknown deviations in instrumental conditions. Fluctuations in retention times across various measurements obscure statistical analysis and thus discovery of relevant patterns in the data[9]. Since retention time shifts are observed in our $^{1}$D chromatogram, we applied the correlation optimized wrapping (COW) algorithm[10] for peak alignment. This method is a piecewise or segmented data preprocessing method (operating on one sample record at a time) aimed at aligning a sample data vector towards a reference vector by allowing limited changes in segment lengths on the sample vector. The outcome of this method contains the correlation between the reference chromatogram retention time and the new chromatogram retention time. With this correlation, a shift time could be found for each peak based on its original retention time and the single peak two-dimensional gas chromatogram could shift on the $^{1}$D based on this shift time.

(5) Summation of individual two-dimensional chromatograms: Adding all individual peaks' two-dimensional chromatograms together after applying the shift time yields the complete two-dimensional chromatogram.

### Section S3: Algorithm description

To distinguish ARDS and non-ARDS patients based on their breath chromatograms, linear discriminant analysis (LDA) was applied to find a linear function that could be used to separate these two groups. However, LDA can only be applied if the number of samples (patient chromatograms) is much larger than the number of features[11] (i.e., the number of VOCs, which was 97 in our study). To overcome this limitation and to decrease the computational complexity of the pattern classifier, principal component analysis (PCA) was applied prior to LDA to reduce the dimensionality of the feature space.

Since PCA is an unsupervised dimension reduction method, it only performs a linear mapping of the data to a lower-dimensional space in such a way that the variance of the data point is maximized. If we directly apply PCA to the overall VOC dataset, the VOCs relevant to ARDS may get overlooked and the interference VOCs that have bigger variance among patients will be kept. Therefore, to produce the best classification result with PCA-LDA, it is critical to find the features (VOCs) that are relevant to ARDS and discard all other interference features. The physical interpretation is that not all of those peaks in the 2D chromatogram may be relevant to ARDS. For example, some of the peaks may be from indoor air background, normal metabolic activities, or other conditions that a patient has. Those irrelevant peaks interfere in the correct classification of ARDS and non-ARDS groups. It is therefore critical to determine which subset of the peaks is most responsible for the differences observed between ARDS and non-ARDS groups.

#### *S3.1. Generation of possible peak subsets*

We first assume that there are a total of *m* different peaks found in all patients' 2D chromatograms with different quantities. For a particular patient not all *m* peaks are present. The quantities of those missing peaks are assigned to be 0. All *m* peaks and their quantities form the entire dataset can be expressed as:

$$\begin{pmatrix} x_{11}, x_{12}, \cdots, x_{1m} \\ x_{21}, x_{22}, \cdots, x_{2m} \\ \vdots \\ x_{p1}, x_{p2}, \cdots, x_{pm} \end{pmatrix}, \quad (1)$$

where $x_{ij}$ is the quantity of the $j^{th}$ peak of the $i^{th}$ patient. In total, there are *m* peaks and *p* patients. We further assume that there are *N* peaks relevant to ARDS and non-ARDS classification.

Consequently, there are $C_m^N$ possible peak combinations (subsets) that can be selected from the dataset in Eq. (1). One such subset can be written as:

$$\begin{pmatrix} x_{1k_1}, x_{1k_2}, \cdots, x_{1k_N} \\ x_{2k_1}, x_{2k_2}, \cdots, x_{2k_N} \\ \vdots \\ x_{pk_1}, x_{pk_2}, \cdots, x_{pk_N} \end{pmatrix}, \quad (2)$$

where $(k_1, k_2, \ldots, k_N)$ is the subset formed by *N* peaks.

#### *S3.2. Criteria of peak subset selection*

PCA was first used for data reduction of the *N* peaks VOC subsets and then LDA was applied to the primary two principal component scores for classification. The total accuracy (true positive plus true negative rate) of classification was used as the criteria for peak subset relevancy to ARDS. For each possible peak subset, PCA was first applied to the *p*-by-*N* dataset to produce *p*-by-*N* principal component scores. Then, the primary two principal component scores (*p*-by-*2*) and the classifier (1 as ARDS and 0 as non-ARDS) for each patient were used to train an LDA model and yield a linear boundary between the ARDS and non-ARDS groups. The total accuracy (number of patients falling in the correct side of the boundary divided by the total patients) was calculated and used as criteria of the relevancy of this VOC subset. Equations (3) and (4) illustrate the methods and processes described above.

$$\begin{pmatrix} x_{1k_1}, x_{1k_2}, \cdots, x_{1k_N} \\ x_{2k_1}, x_{2k_2}, \cdots, x_{2k_N} \\ \vdots \\ x_{pk_1}, x_{pk_2}, \cdots, x_{pk_N} \end{pmatrix} \xRightarrow{PCA} \begin{pmatrix} s_{11}, s_{12}, \cdots, s_{1N} \\ s_{21}, s_{22}, \cdots, s_{2N} \\ \vdots \\ s_{p1}, s_{p2}, \cdots, s_{pN} \end{pmatrix}, \quad (3)$$

where $s_{ij}$ is the $j^{th}$ principal component score of the $i^{th}$ patient.

$$\begin{pmatrix} s_{11}, & s_{12} \\ s_{21}, & s_{22} \\ \vdots & \\ s_{p1}, & s_{p2} \end{pmatrix} with \begin{pmatrix} c_1 \\ c_2 \\ \vdots \\ c_p \end{pmatrix} \xRightarrow{LDA} linear\ boundary \xrightarrow{yields} Classification\ accuracy, \quad (4)$$

where $c_i$ is the classifier (1 for ARDS and 0 for non-ARDS) of the $i^{th}$ patient. Finally, the peak combinations (i.e., subsets) with highest accuracy were selected. For each of these selected peak combinations (subsets), the patients' coordinates on the PCA plot were decided by their principal component scores. The mean distance of the 20% patients closest to the boundary line, normalized by the mean distance, was calculated. The peak subset with highest boundary distance was chosen as the optimal peak subset.

#### *S3.3. Iterative peak subset selection*

Since human breath is a complex mixture, the total peak number $m$ is large and the total number of possible combinations of peaks (i.e., peak subsets), $\sum_{N=1}^{m} C_m^N$ , is enormous. As a result, it requires a great amount of computation time to evaluate all the peak subsets. To expedite the selection process, in this work we started with peak subsets formed by a small number of peaks (e.g., $n$ peaks, which resulted in $C_m^n$ subsets to be evaluated). Once the most relevant peak subset was determined, more peaks were added to this selected subset, aiming to achieve higher accuracy.

Assuming that there are *n'* more peaks that are relevant to ARDS (*n'* is another small number of VOCs in order to save the computation time), $C_{m-n}^{n'}$ possible peak combinations (subsets) can be found and added to the previously optimized VOC subset to form a new peak subset, i.e.,

$$\begin{pmatrix} x_{1k_1}, x_{1k_2}, \cdots, x_{1k_n}, x_{1l_1}, x_{1l_2}, \cdots, x_{1l_{n'}} \\ x_{2k_1}, x_{2k_2}, \cdots, x_{2k_n}, x_{2l_2}, x_{2l_2}, \cdots, x_{2l_{n'}} \\ \vdots \\ x_{pk_1}, x_{pk_2}, \cdots, x_{pk_n}, x_{pl_2}, x_{pl_2}, \cdots, x_{pl_{n'}} \end{pmatrix}, \quad (5)$$

where $(l_1, l_2, \ldots, l_{n'})$ is the peak subset with $n'$ peaks.

With the new peak subset, PCA and LDA were applied to calculate the accuracy and the boundary distance. If the classification accuracy increased or the boundary distance increased, those *n'* peaks were kept and more peaks out of *m-n-n'* peaks would be added iteratively in the same manner described above. If accuracy no longer increased or the boundary distance no longer increased, then the iteration process was ended, and the final optimal peak subset was determined. The overall iteration processes are illustrated in Figure S2.

#### *S3.4. Training and testing with ARDS and non-ARDS patients*

The entire patient data set was divided into two sets: training set (*p* patients) and testing set (*q* patients). The training set was used to select the optimal peak subset for best classification and determine the linear boundary, whereas the testing set was used to validate the determined optimal peak subset and the boundary on the PCA plot.

Assuming the final optimal peak subset containing *N* peaks. With the *N* peaks subset, the PCA analysis yields an *N*-by-*N* PCA coefficient and a linear boundary line between ARDS and non-ARDS groups.

$$\begin{pmatrix} x_{t_1k_1}, x_{t_1k_2}, \cdots, x_{t_1k_N} \\ x_{t_2k_1}, x_{t_2k_2}, \cdots, x_{t_2k_N} \\ \vdots \\ x_{t_pk_1}, x_{t_pk_2}, \cdots, x_{t_pk_N} \end{pmatrix} \overset{PCA}{\Longrightarrow} \begin{pmatrix} Coeff_{11}, Coeff_{12} \dots, Coeff_{1N} \\ Coeff_{21}, Coeff_{22} \dots, Coeff_{2N} \\ \vdots \\ Coeff_{N1}, Coeff_{N2} \dots, Coeff_{NN} \end{pmatrix} \text{ and } \begin{pmatrix} s_{t_1 1}, s_{t_1 2} \dots, s_{t_1 N} \\ s_{t_2 1}, s_{t_2 2} \dots, s_{t_2 N} \\ \vdots \\ s_{t_p 1}, s_{t_p 2} \dots, s_{t_p N} \end{pmatrix}, \quad (6)$$

where $(t_1, t_2, \dots t_p)$ is the training set patients; in total there are $p$ patients in the training set.

$$\begin{pmatrix} s_{t_1 1}, s_{t_1 2} \\ s_{t_2 1}, s_{t_2 2} \\ \vdots \\ s_{t_p 1}, s_{t_p 2} \end{pmatrix} with \begin{pmatrix} c_{t_1} \\ c_{t_2} \\ \vdots \\ c_{t_p} \end{pmatrix} \overset{LDA}{\Longrightarrow} linear\ boundary \quad (7)$$

With the N-by-N PCA coefficient acquired from the training set, the PC scores of the testing set can be calculated by multiplying the PCA coefficient to the N peak subset of all testing patients. With the linear boundary line acquired from the training set, the final classification accuracy can be calculated.

$$\begin{pmatrix} x_{v_1k_1}, x_{v_1k_2}, \cdots, x_{v_1k_N} \\ x_{v_2k_1}, x_{v_2k_2}, \cdots, x_{v_2k_N} \\ \vdots \\ x_{v_qk_1}, x_{v_qk_2}, \cdots, x_{v_qk_N} \end{pmatrix} \cdot \begin{pmatrix} Coeff_{11}, Coeff_{12} \dots, Coeff_{1N} \\ Coeff_{21}, Coeff_{22} \dots, Coeff_{2N} \\ \vdots \\ Coeff_{N1}, Coeff_{N2} \dots, Coeff_{NN} \end{pmatrix} \xrightarrow{yields} \begin{pmatrix} s_{v_1 1}, s_{v_1 2}, \cdots, s_{v_1 N} \\ s_{v_2 1}, s_{v_2 2} \dots, s_{v_2 N} \\ \vdots \\ s_{v_q 1}, s_{v_q 2} \cdots, s_{v_q N} \end{pmatrix}, \quad (8)$$

where $(v_1, v_2, \dots, v_q)$ is the testing set patients; in total there are $q$ patients in testing set;

$$\begin{pmatrix} s_{v_1 1}, s_{v_1 2} \\ s_{v_2 1}, s_{v_2 2} \\ \vdots \\ s_{v_q 1}, s_{v_q 2} \end{pmatrix} \text{with } linear\ boundary \xrightarrow{yields} final\ classification\ accuracy \quad (9)$$

#### *S3.5. Selection of the optimal subset of peaks relevant to ARDS*

In our study, a total of *m=97* peaks were found in 2D chromatograms. We first assumed that there are n=4 peaks relevant to classification of ARDS and non-ARDS. We found that the 4-peak subset of Peak #(2, 44, 72, 79) provides the best classification with the total accuracy of 88.4% (see the corresponding peaks on the 2D GC chromatogram in Figure S3 and the PCA-LDA results in Figure S4(a)). Then n'=5 peaks were added and we found that the 9-peak subset of [(2, 44, 72, 79) + (34, 38, 62, 66, 81)] provides the best classification with the total accuracy of 93.0% (see the corresponding peaks on a 2D GC chromatogram in Figure S3 and the PCA-LDA results in Figure S4(b)). Then another n'=5 peaks were added and we found that the 14-peak subset of [(2, 44, 72, 79) + (34, 38, 62, 66, 81) + (54, 61, 63, 71, 75)] provides the best classification with the total accuracy of 93.0% (see the corresponding peaks on a 2D GC chromatogram in Figure S3 and the PCA-LDA results in Figure S4(c)). Since the classification accuracy and the boundary distance does not improve from the 9-peak subset to the 14-peak subset (i.e., the ARDS and non-ARDS groups are not clustered/separated better), the 9-peak subset was selected as the final optimal peak subset, which consists of Peak #(2, 44, 72, 79, 34, 38, 62, 66, 81) in the 2D chromatogram.

### Section S4: Comparison of portable 2D GC and other technologies

Table S1 lists the peak capacity estimated from three exemplary peaks. For GC × GC, the peak capacity is defined as $n_{GC\times GC} = n_1 \times n_2$, where $n_1$ and $n_2$ are the peak capacity for $^1$D and $^2$D, respectively[1]. The conventional method for calculation of peak capacity using 4σ - bottom-to-bottom-width $w_{4\sigma}$ is given by: $n_{4\sigma} = (t_R^1/w_{4\sigma}^1)(CP_M/w_{4\sigma}^2)$, where C is the number of channels in $^2$D and $P_M$ is the modulation period. The peak capacity value for three selected peaks is listed in the table below as $n_{4\sigma}$.

SIFT-MS and IMS technologies use the full-width-at-half-maximum (FWHM, $w_{FWHM} = 0.588 w_{4\sigma}$) to evaluate their resolution (or resolving power)[12-14]. To compare our results with their resolving power we calculate the 2D GC peak capacity using $w_{FWHM}$, i.e., $n_{FWHM} = (t_R^1/w_{FWHM}^1)(CP_M/w_{FWHM}^2)$. The peak capacity value for the three selected peaks is listed in the table below as $n_{FWHM}$. The peak capacity of the $^1$D separation ranges 50-100. The 2D peak capacity ranges 600-1,200, which is much larger than the resolution of 10-30 in IMS[15,16] (note: some high-resolution IMS can have a resolution over 100) and of 200-300 in SIFT-MS[17-19].

| Peak # | $t_R^1$ | $w_{FWHM}^1$ | $w_{4\sigma}^1$ | $t_R^2$ | $w_{FWHM}^2$ | $w_{4\sigma}^2$ | $n_{FWHM}$ | $n_{4\sigma}$ |
|---|---|---|---|---|---|---|---|---|
| 10 | 154.0 s | 2.8 s | 4.8 s | 5.0 s | 0.9 s | 1.5 s | 1,222 | 423 |
| 19 | 315.7 s | 3.9 s | 6.6 s | 6.0 s | 1.5 s | 2.5 s | 1,079 | 373 |
| 30 | 544.3 s | 4.1 s | 7.0 s | 10.4 s | 4.0 s | 6.8 s | 664 | 230 |

Table S1. Peak capacity for the portable 2D GC calculated for three exemplary peaks.

$t_R^1$: $^1$D retention time

$t_R^2$: $^2$D -dimensional retention time

$w_{4\sigma}^1$: $^1$D peak width (4σ - bottom-to-bottom)

$w_{4\sigma}^2$: $^2$D peak width (4σ - bottom-to-bottom)

$w_{FWHM}^1$: $^1$D peak width (FWHM)

$w_{FWHM}^2$: $^2$D peak width (FWHM)

C: Number of $^2$D channels (in our case: C = 2)

$P_M$: modulation time (in our case: $P_M$ = 10 s)

FWHM: Full width at half maximum

MS: Spectrometry

IMS: Ion mobility spectrometry

SIFT: Selected-ion flow-tube mass spectrometry

**Section S5: Patient medical history during time series testing dates**

Patient #2 was a healthy subject tested for 4 days.

Patient #3 was a healthy subject tested for 4 days.

Patient #7 was sampled for 4 days and had ARDS by the Berlin Criteria since the 1st testing day. No signs of recovery at least 4 days after the last testing day.

Patient #11 was a potential and undetermined ARDS patient on the 1st test day and then upgraded to ARDS on the next day. By the Berlin Criteria he/has ARDS for all 3 days.

Patient #12 was suspected for pneumonia on the 1st testing day. This patient was tested for 3 days and no ARDS was developed during this period based on the Berlin Criteria.

Patient #27 was a potential and undetermined ARDS patient on the 1st test day and then upgraded to ARDS on the next day. Based on the Berlin Criteria he/has ARDS for all 3 days.

Patient #30 had pneumonia and ARDS based on the Berlin Criteria since 1st test day and was tested for 3 days. No signs of recovery and was shifted to comfort care after the last testing day.

Patient #31 had acute respiratory failure on the 1st testing day but no ARDS based on the Berlin Criteria during the 2 testing days.

Patient #34 had hypoxemic respiratory failure on the 1st testing day but no ARDS based on the Berlin Criteria during the 3 testing days.

Patient #35 had no ARDS based on the Berlin Criteria during the 2 testing days.

Patient #36 had ARDS since the 1st sampling day. On the 3rd day the patient was still listed as ARDS patients based on the Berlin Criteria and then got extubated and discharged from ICU on the 5th day.

Patient #38 was a healthy subject tested for 3 days.

Patient #39 had ARDS based on the Berlin Criteria during the 2 testing days. No signs of recovery.

Patient #40 had pneumonia on the 1st testing day but no ARDS based on the Berlin Criteria during the 2 testing days.

Patient #42 had ARDS based on the Berlin Criteria during the 2 testing days. No signs of recovery.

Patient #45 had ARDS based on the Berlin Criteria during the 5 testing days. No signs of recovery and was shifted to comfort care on the last testing day.

Patient #46 had no ARDS based on the Berlin Criteria during the 3 testing days.

Patient #47 was sampled for 4 days and got extubated and discharged on the 6th day. Based on the Berlin Criteria this patient had ARDS for all first 4 days.

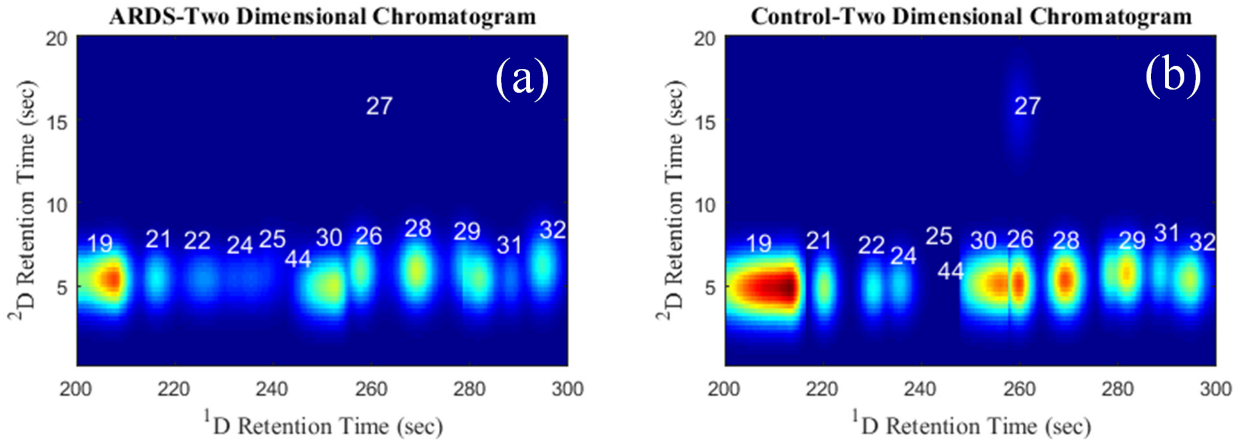

Figure S1. (a) Zoomed-in portion of Figure 4(b). (b) Zoomed-in portion of Figure 4(b).

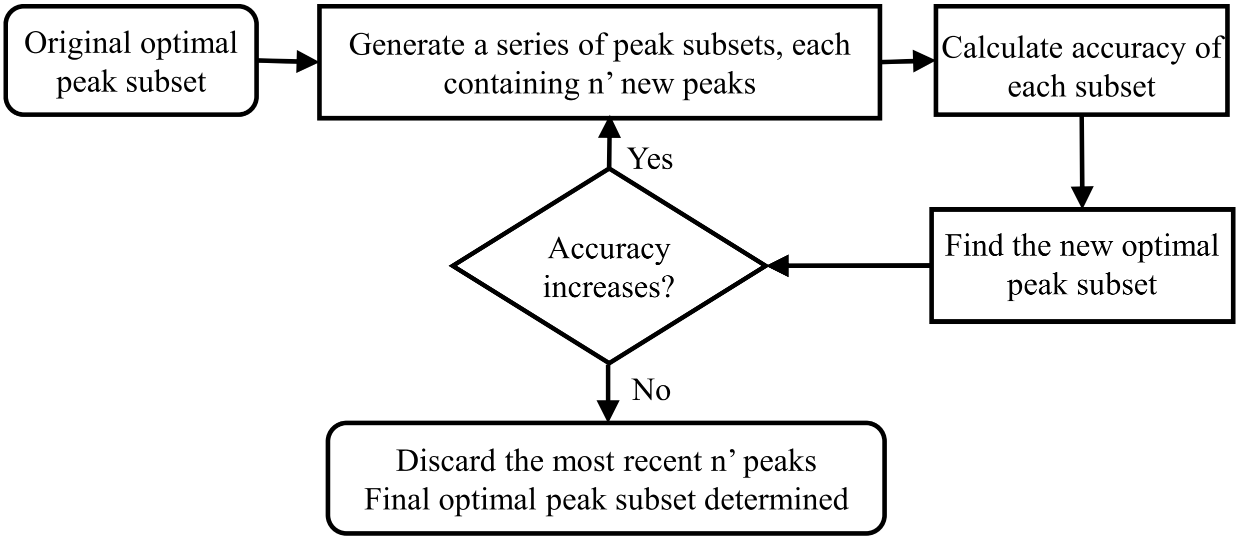

Figure S2. Iterative peak subset selection procedure.

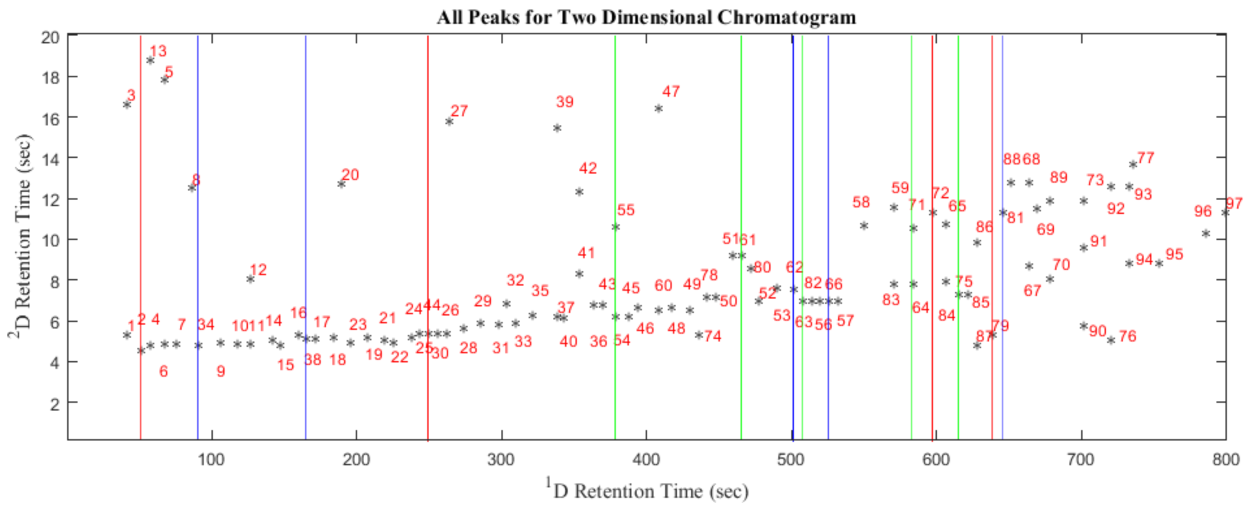

Figure S3. Selection of the optimal subset of peaks relevant to ARDS. Red lines mark the $^{1}$D retention time of the 4 peaks selected in the first iteration. Blue lines mark the $^{1}$D retention time of the additional 5 peaks selected in the second iteration. Green lines mark the $^{1}$D retention time of the additional 5 peaks selected in the third iteration. Peak #34 in the 9-peak subset nearly co-elutes with Peak #8. Peak #54 and #71 in the 14-peak subset co-elutes with Peak #55 and Peak #64, respectively.

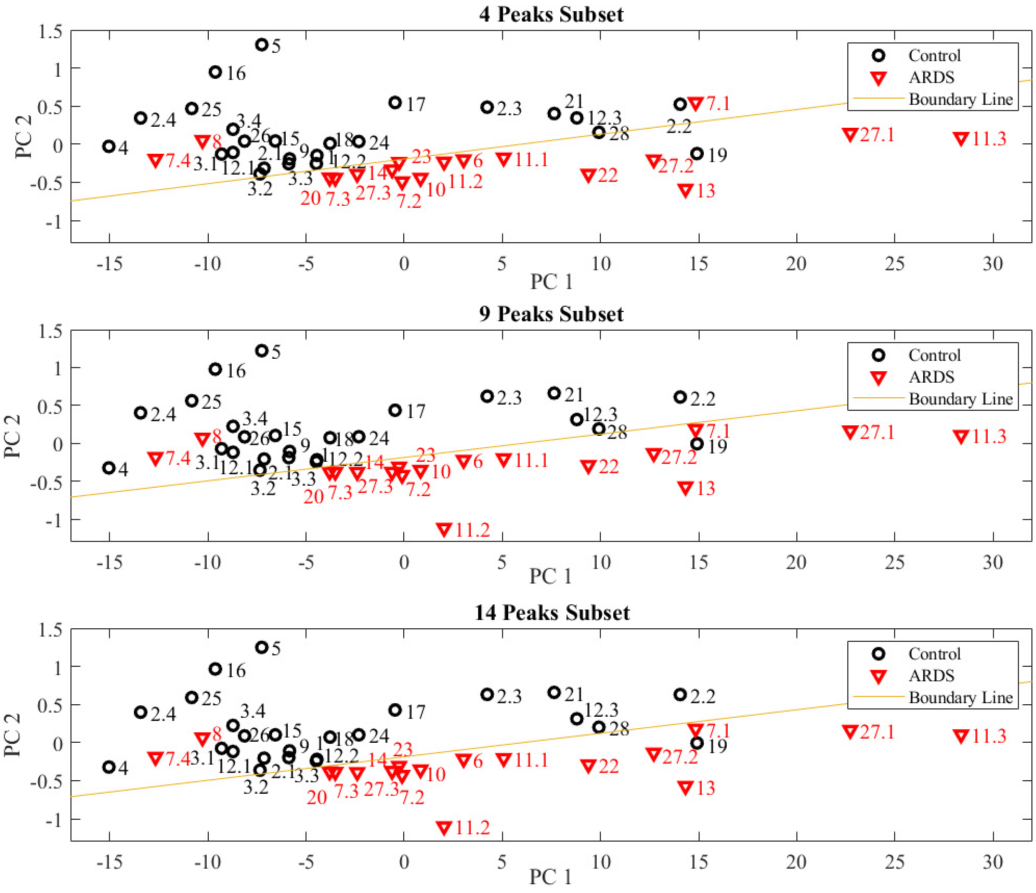

Figure S4. PCA plots using the subset containing 4 peaks, 9 peaks, and 14 peaks for the training set of patients. The red and black symbols denote respectively the ARDS and non-ARDS patients adjudicated by physicians using the Berlin criteria. The patient numbers are given by the symbol. For example, “11.1” and “11.3” denote Patient #11, Day 1 and Day 3 results, respectively. The bottom/top zone below/above the boundary line represents respectively the ARDS/non-ARDS region using the breath analysis method.

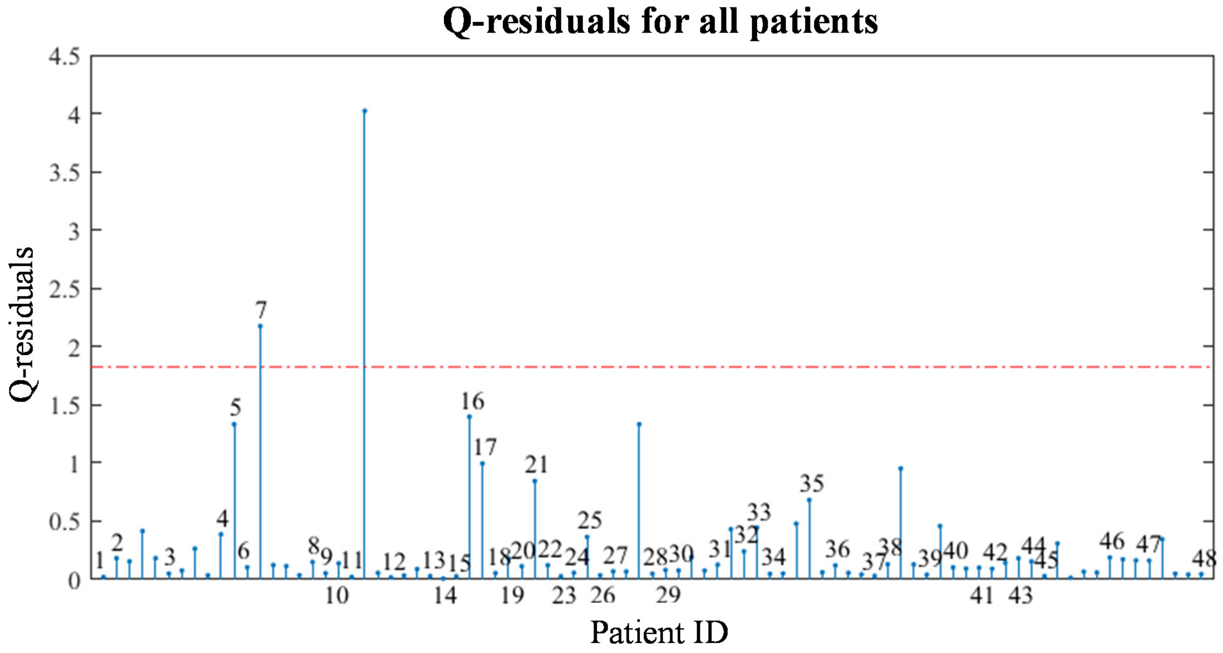

Figure S5. Q-residuals of the PCA model (Figure 7) for all recruited patients. For the patients with time series tests, only the 1st test day is marked with the patient ID. The red dashed curve shows the 99% confidence level.

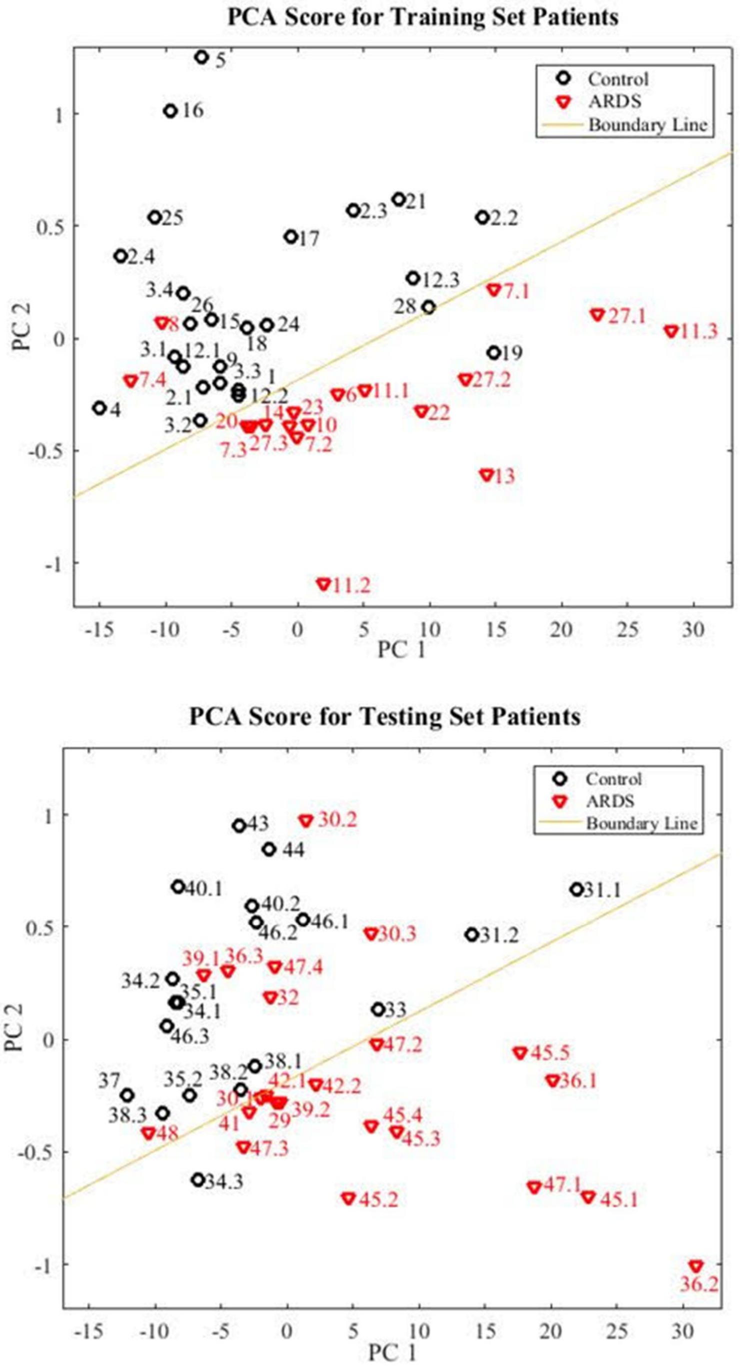

Figure S6. PCA plot for the training and testing set of patients. The corresponding statistics is given in Table S2.

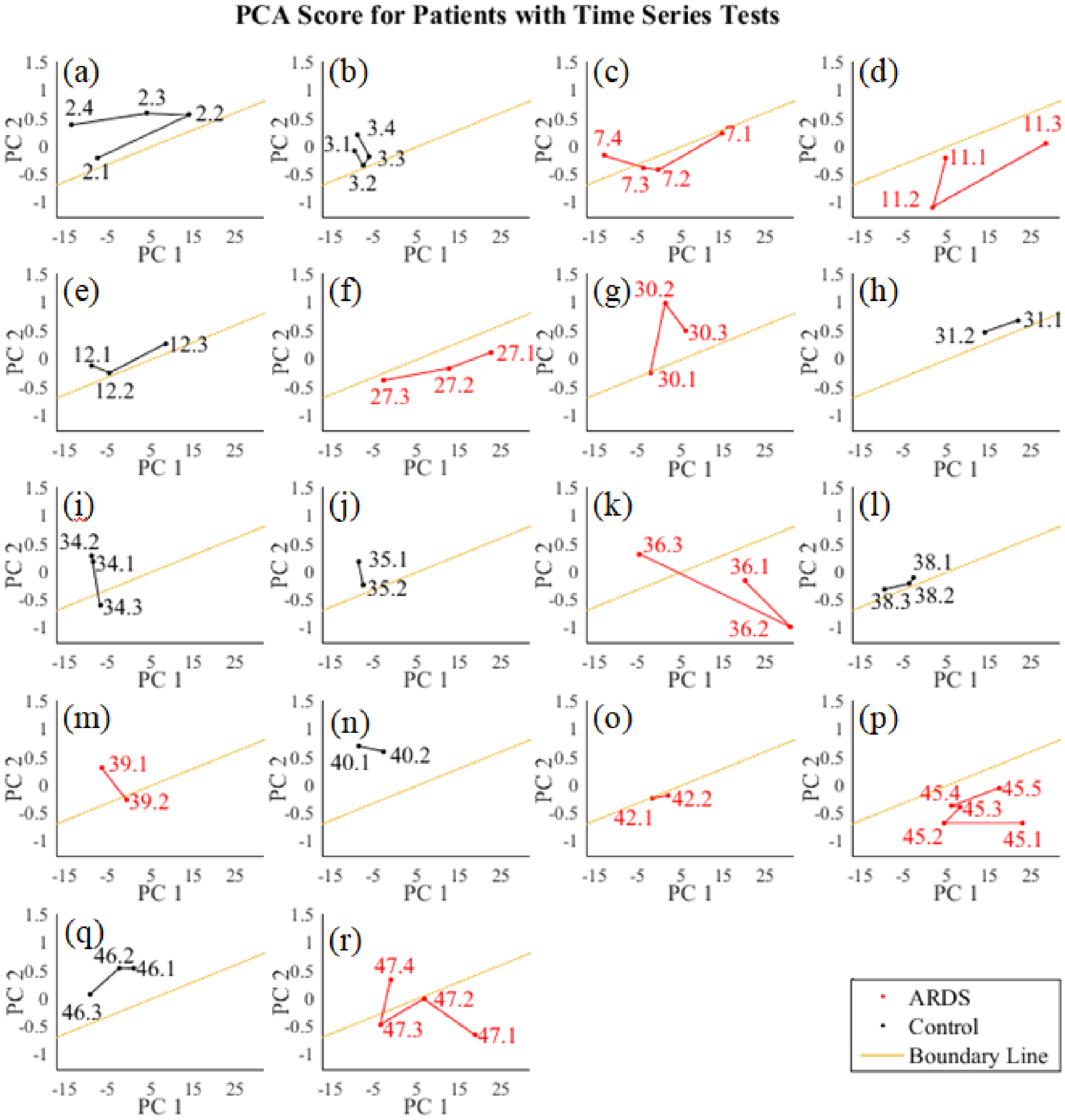

Figure S7. Individual trajectories of all 18 patients with time series tests on the PCA plot. Refer to Section S5 for the patient medical history.

| Training Set | **ARDS** | **Non-ARDS** | **Total** |
|---|---|---|---|
| **Positive (our results)** | 16 | 1 | 17 |
| **Negative (our results)** | 2 | 24 | 26 |
| **Column total** | 18 | 25 | 43 |
| **Specificity** | 1-1/25=96.0% | | |
| **Sensitivity** | 1-2/18=88.9 % | | |
| **Positive predictive value** | 16/17=94.1% | | |
| **Negative predictive value** | 24/26=92.3% | | |
| **Total accuracy** | 40/43=93.0% | | |
| Testing Set | **ARDS** | **Non-ARDS** | **Total** |
| **Positive (our results)** | 16 | 1 | 17 |
| **Negative (our results)** | 7 | 18 | 25 |
| **Column total** | 23 | 19 | 42 |
| **Specificity** | 1-1/19=94.7 % | | |
| **Sensitivity** | 1-7/23=69.6 % | | |
| **Positive predictive value** | 16/17=94.1% | | |
| **Negative predictive value** | 18/25=72.0% | | |
| **Total accuracy** | 34/42=80.9% | | |

Table S2. Statistics for the training and testing set of patients.
